# Spatial inheritance patterns across maize ears are associated with alleles that reduce pollen fitness

**DOI:** 10.1101/2025.09.17.676879

**Authors:** Diana Ruggiero, Michelle Bang, Marilyn Leary, Harrison Flieg, Luis Garcia-Lamas, Zuzana Vejlupkova, Molly Megraw, Duo Jiang, Samuel Leiboff, John E Fowler

**Affiliations:** Oregon State University Department of Botany & Plant Pathology; Oregon State University Department of Statistics

**Keywords:** *Zea mays* subsp. *mays* L., high-throughput phenotyping, pollen fitness, pollination, plant reproduction, computer vision, genetics, Faster R-CNN, seed, inflorescence

## Abstract

Early studies noting uneven spatial distribution of progeny genotypes after pollination support a hypothesis where differences in pollen tube growth rate can bias inheritance. We used computer vision and statistical analysis to show alleles reducing maize pollen fitness are likely to produce statistically significant increasing, decreasing, or curvilinear spatial patterns from the apex of the inflorescence to the base, suggesting that differential pollen tube growth is not the only mechanism at play.

**Summary:** Often, more pollen grains land on recipient flowers than there are ovules to fertilize. Consequently, the haploid male gametophyte engages in post-pollination competition, one way that pollen genotype can influence inheritance. The maize (*Zea mays* subsp. *mays* L.) inflorescence (ear), with its elongated stigma and style structures (silks), has a conspicuous spatial heterogeneity, with longer silks at the base of the ear than those at the apex. To evaluate the hypothesis that alleles with reduced pollen fitness influence the spatial distribution of progeny genotypes along the ear, we developed an updated phenotyping platform that maps mutant *Ds-GFP* kernel phenotypes on the ear via an implementation of the Faster R-CNN machine vision model (EarVision.v2) and a statistical pipeline that evaluates the relationship between kernel position and transmission ratio (EarScape). In our dataset (1384 ears) representing 58 *Ds-GFP* alleles, none with Mendelian inheritance (0/48) showed any significant pollen-conditioned spatial trend. In contrast, 50% of alleles with a pollen-specific transmission defect (5/10) exhibited significant spatial effects. An insertion into a gene encoding a putative actin-binding protein, *base-to-apex gradient1\** (*bag1\**), conditions increased mutant transmission at the ear apex relative to the base. Surprisingly, mutant alleles of two other pollen-expressed genes can generate the opposite pattern, decreased mutant transmission toward the ear apex; and two mutant alleles of the sperm-cell attachment factor, *gamete expressed2* (*gex2*), can produce ears with transmission highest at both base and apex. We conclude that pollen fitness mutants have relatively common but heterogenous effects on the spatial distribution of progeny genotypes.

## Introduction

In flowering plants, the alternation of generations leads to unique dynamics in gamete competition. As in animals, the competitive objective is the same: the first sperm cell to reach the egg contributes alleles to the progeny. However, unlike animals, delivery of the sperm cells to the ovule requires the gametophyte (pollen grain and pollen tube), which provides transport to the stigma and then navigates through the style to the embryo sac. Also unlike animal sperm, pollen is transcriptionally active, and its haploid genome directly drives gene expression both during its development and during pollen tube growth (Qin *et al*., 2009; Johnson *et al*., 2019; Nelms and Walbot, 2022; Somers and Nelms, 2023; Misra *et al*., 2025). Because pollen is haploid, loss-of-function alleles are exposed and this direct phenotypic expression provides potential for selection at the gametophytic stage (Mulcahy and Mulcahy, 1987). Successful fertilization and allele transmission thus involves a complex interaction of the male gametophyte (and its genotype) with the carpel tissue and embryo sac (Johnson *et al*., 2019).

One way that pollen genotype influences allele transmission is through competition between pollen tubes within the carpel. More pollen typically lands on the stigma than there are ovules to fertilize, resulting in post-pollination competition (Stephenson and Bertin, 1983). As multiple pollen tubes race to the ovule through the style, competition occurs both between pollen from different individuals and among pollen from the same parent (Heslop-Harrison *et al*., 1985; Birkhead and Møller, 1998; Swanson *et al*., 2016). Long-standing observations highlight the importance of gametophytic traits, especially pollen tube growth rate, and how these traits could interact with spatial features of the female reproductive structure, such as the distribution of ovules along a transmitting tract (Brink and MacGillivray, 1924; Walsh and Charlesworth, 1992). Pollen genotype could therefore influence success of fertilization, producing spatial patterns of genotype frequencies in the resulting progeny (Hill and Lord, 1986; Quesada *et al*., 1991).

Spatial patterns in fertilization, influenced by pollen genotype, have been observed in unmanipulated flowers from taxa with polyovulate ovaries, including *Raphanus* (Marshall and Ellstrand, 1986; Hill and Lord, 1986; Marshall, 1991; Marshall and Diggle, 2001; Marshall and Evans, 2016), *Cucurbita* (Quesada *et al*., 1991), and the legumes (Cooper and Brink, 1940; Barnes and Cleveland, 1963; Ibarra-Perez *et al*., 1996). These studies reveal positional effects on seed abortion, allele transmission, siring success, and ovule fertilization order, although the genetic bases of these effects have not been determined. Pollen tube growth rate has been proposed as a key determinant of competitive success (Correns, 1928; Walsh and Charlesworth, 1992; Williams and Reese, 2019), and manipulating the distance the pollen tube must travel to the ovary impacts competition and resulting fertilization (Brink, 1925; Mulcahy and Mulcahy, 1975; Pélabon *et al*., 2016). For example, Brink (1925) observed reduced transmission of the *waxy* pollen mutant in maize when fertilization occurred through longer styles. In *Arabidopsis thaliana*, biased inheritance across the silique (fruit) was observed at positions farther removed from the stigma for mutations that reduce pollen fitness (Meinke, 1982; Goubet *et al*., 2003; Cole *et al*., 2005).

Given the ease of collecting large quantities of pollen for controlled pollinations, maize is an attractive model for studying pollen function. Maize pollen is also notable for extremely rapid germination and pollen tube growth. A pollen grain landing on the silk, the elongated stigma that also functions as the style, germinates within 5–10 minutes (Heslop-Harrison *et al*., 1984; Dresselhaus *et al*., 2011). The germinated pollen tube then grows into the silk cortex to reach one of two transmitting tracts (Figure 1b; Dresselhaus *et al*., 2011). Within the silk, pollen tubes can grow at rates exceeding 1 cm/hr, typically achieving fertilization within 16–24 hours post-pollination (Bedinger, 1992; Mascarenhas, 1993; Sheridan and Clark, 1994). Maize may possess the longest transmitting tract among angiosperms and exhibits pollen tube growth rates up to 100 times faster than *Arabidopsis*—features that may contribute to a substantial role for pollen competition in its evolutionary history (reviewed in Williams and Reese, 2019).

**Figure 1:**
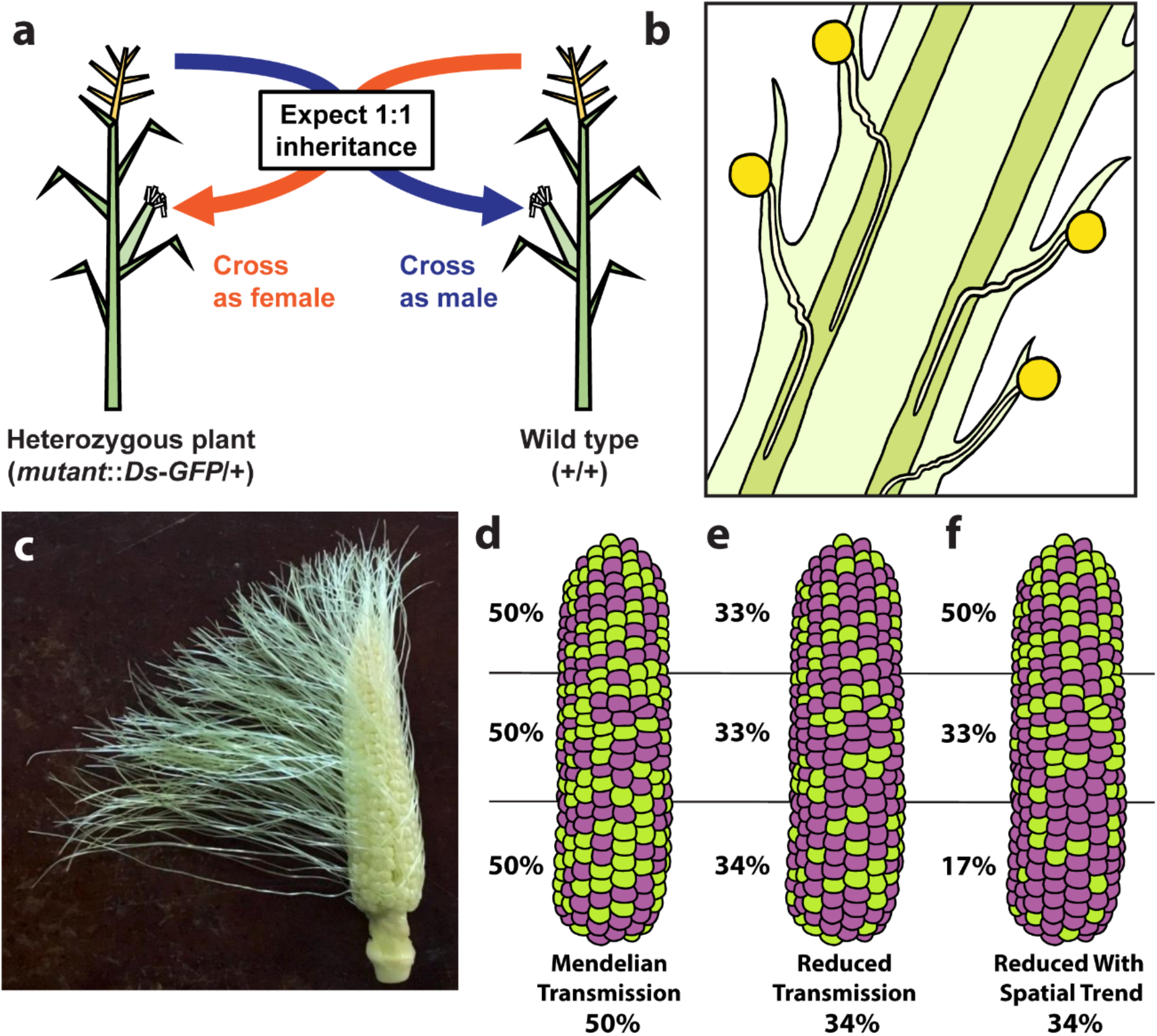
Observing pollen fitness effects and non-random kernel distribution on the maize ear. (a) Reciprocal outcrosses between a maize plant heterozygous for a *Ds-GFP* insertion allele (*mutant::Ds-GFP*/+) and a wild type plant (+/+) both as a male (using the pollen) and as a female (using the ear, control). 1:1 Mendelian transmission in the pollen cross is expected unless an allele is associated with reduced pollen fitness and a resulting transmission defect. (b) Multiple maize pollen tubes grow down the transmitting tracts of the silk, leading to competition among male gametophytes. (c) After trimming in preparation for controlled pollination, the silks at the apex of the ear are shorter than the silks at the base, generating a gradient in the distance required for pollen tubes to reach each embryo sac. (d-f) Three different hypothetical transmission patterns of the *Ds-GFP* allele (green kernels) and wild type allele (purple kernels). (d) No spatial pattern, Mendelian transmission of *Ds-GFP* allele with equal numbers of fluorescent and non-fluorescent kernels, equal distribution along the length of the ear. (e) No spatial pattern, reduced transmission of *Ds-GFP* allele, equal distribution along the length of the ear. (f) Increasing linear pattern toward the apex, reduced transmission of *Ds-GFP* allele (33.6% TR), unequal distribution along the length of the ear.

The maize inflorescence (the ear) also exhibits pronounced spatial heterogeneity. Due to the vertical arrangement of florets along the ear, silks associated with ovules at the base are longer than those at the apex—an effect heightened when silks are trimmed before a controlled pollination (Figure 1c). Silk emergence and growth follows an apical-basial gradient, likely related to the sequence of ovule maturation; the earliest-emerging silks are associated with ovules located near the base of the ear, with silks closer to the apex emerging later (Oury *et al*., 2016; Oury *et al*., 2022). Because each silk is associated with a single ovule, maize exhibits a high pollen-to-ovule ratio, suggesting high pollen competition (Erbar, 2003). Upwards of 30 pollen tubes can grow simultaneously in a single maize silk (Heslop-Harrison *et al*., 1985), providing ample opportunity for pollen genotype to influence success of fertilization across silks of different length and maturity.

The long history of genetic studies in maize has identified loci that influence male gametophyte function, often using linked endosperm markers to screen for transmission rate distortion (Mangelsdorf and Jones, 1926). Recently, this approach has been greatly expanded using insertion alleles from the Dooner/Du collection of transposable element lines (acdsinsertions.org), which are marked with an endosperm-expressed fluorescent GFP reporter (Li *et al*., 2013; Warman *et al*., 2020). Conceptually, controlled pollinations between known parental genotypes should generate ears with distinctive progeny distributions across the ear, visualized using the *Ds-GFP* marker (Figure 1a, d-f). Mendelian transmission (Figure 1d) can be distinguished from globally reduced transmission due to a pollen fitness defect (Figure 1e), and from spatially biased patterns of inheritance (Figure 1f). The large size of each ear, the high numbers of progeny resulting from a single pollination, and the ability to track inheritance easily with endosperm markers make maize an experimental model well-suited to investigating the full range of possible pollen-driven spatial inheritance patterns.

Quantifying spatial bias in maize fertilization depends on phenotyping ears and kernels at scale while processing positional information. Advances in machine learning and computer vision have increased the feasibility and effectiveness of high-throughput phenotyping approaches, enhancing the potential for identifying subtle phenotypes (Warman and Fowler, 2021). A range of platforms for imaging and analyzing maize ears and kernels have been developed (Miller *et al*., 2017; Adke *et al*., 2021; Gonzalez *et al*., 2022; Shi *et al*., 2022), including several with rotating scanners that generate projected images of the entire circumference of an ear (Zhang *et al*., 2020; Warman *et al*., 2021; Gillette *et al*., 2023), a critical feature for determining kernel position. For example, one such system enabled detection of spatially distinctive patterns of kernel abortion along the ear axis in response to stress (Oury *et al*., 2022).

Building on these advances, we developed an improved phenotyping pipeline (EarVision.v2) to test the hypothesis that alleles with reduced pollen fitness impact the spatial distribution of progeny kernels on the ear. By integrating automated phenotyping with a statistical modeling pipeline (EarScape), we assessed fertilization patterns of hundreds of ears comprising thousands of progeny kernels, producing a comprehensive assessment of 58 individual alleles for spatial transmission effects. Among the ten alleles with reduced pollen fitness, five also showed significant spatial bias in transmission along the ear. Surprisingly, these five alleles exhibited diverse spatial transmission trends, encompassing ears showing increasing, decreasing, and curvilinear patterns. These trends are inconsistent with a model where spatial differences are driven solely by defective pollen tube growth in the insertion mutants (i.e., only resulting in patterns of reduced transmission toward the ear base).

## Results

### An improved ear and kernel phenotyping platform supports large-scale assessment of potential spatial inheritance effects

To assess the relationship between allele transmission and spatial patterns of inheritance across the maize inflorescence, we sought to analyze the broadest possible range of potential pollen fitness mutants, using ear and kernel phenotyping from five years of heterozygous outcrosses with *Ds-GFP*-marked alleles (Figure 1a). To accomplish this comprehensive assessment, we developed an improved iteration of our phenotyping platform that assesses the transmission ratios and spatial trends of maize alleles (MES.v2 and EarVision.v2) by automatically detecting and mapping both fluorescent and non-fluorescent kernels in scanned ear images. These build upon the initial version of the platform (Warman *et al*., 2021) for *Ds-GFP* fluorescent kernel detection, featuring a redesigned ear scanner, a reimplemented object detection system, enhanced internal quality control, and, critically, reporting of kernel position on the ear base-to-apex axis, which makes analysis of spatial patterns possible.

To increase throughput, the Maize Ear Scanner (MES.v2; Figure 2a,b) was reimplemented on a lightweight, compact aluminum chassis, with a benchtop footprint of 20 x 50 cm (Appendix S1). This chassis creates a stable rail system for the mounted camera, reducing the potential for camera movement between shots. Rotation is controlled by a stepper motor driven by an Arduino microcontroller, which provides a smooth, controlled spin for imaging, as well as programmatic control of the rotational speed. MES.v2 uses an adjustable set of paddles with 3D printed adapters to load and unload maize ears, reducing the time spent preparing each ear for imaging. Overall, the improvements allow for an ear to be loaded and imaged in approximately half the time of the MES.v1 scanner. As with the original system, the scanner camera captures a video of the full ear circumference, which is then projected into a 2D image (Figure 2c-f). To capture kernel phenotypes using full-spectrum visible light (e.g., for anthocyanin pigment), we implemented softbox lighting (Figure 2c-g), whereas a blue light and an orange camera filter enable photography of dose-dependent GFP kernel phenotypes (Figure 2d-h).

**Figure 2:**
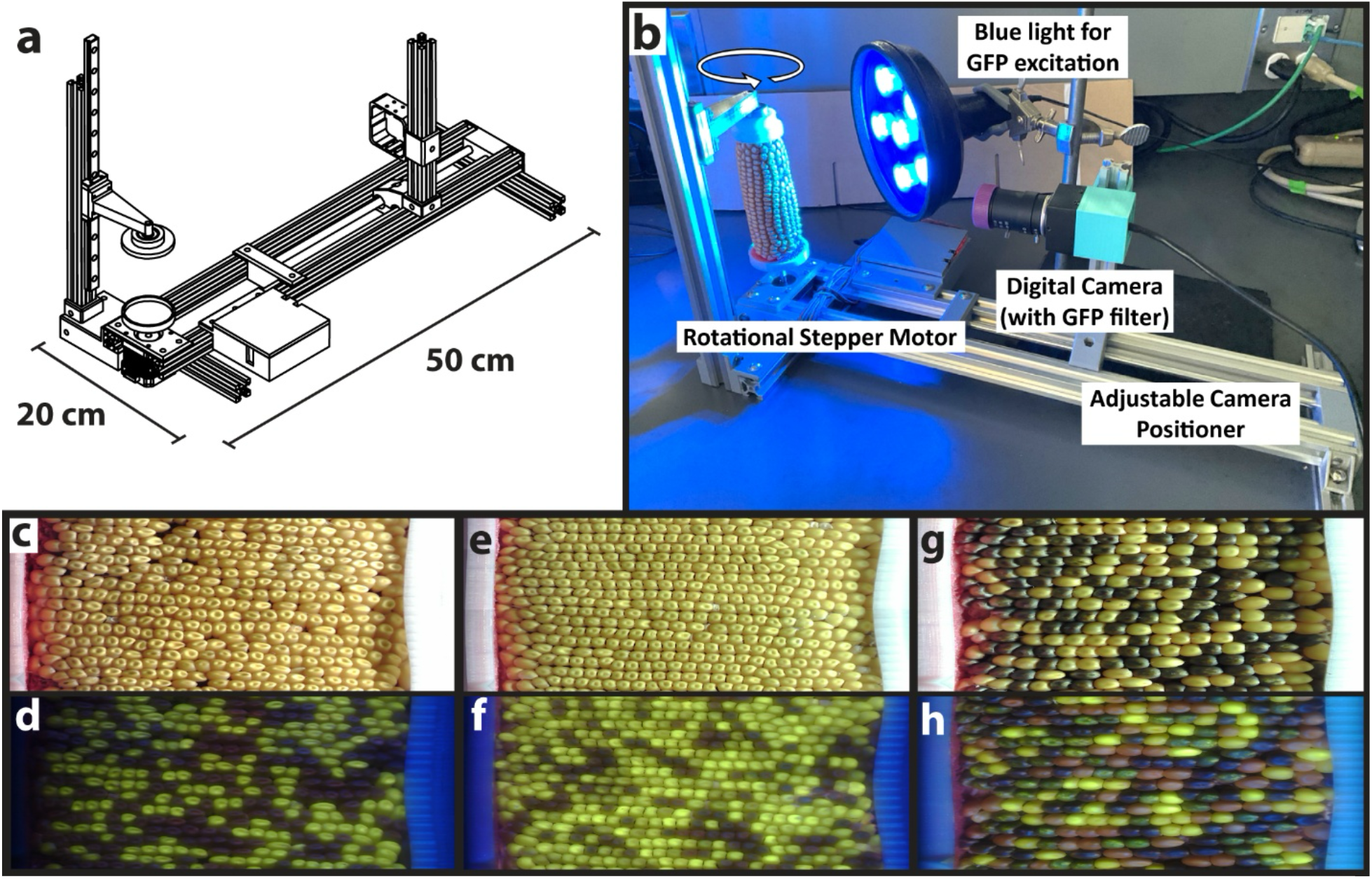
The compact MES.v2 ear scanner efficiently produces images enabling assessment of *Ds-GFP* allele inheritance. (a) Design schematic of MES.v2 scanner assembly. (b) Scanner in operation with camera, ear, and blue light for GFP excitation. (c-h) 2D image projections generated from rotational videos collected by MES.v2. (c,e,g) Full-spectrum visible light images of ears. (d,f,h) Blue light/orange filter images of ears showing fluorescent *Ds-GFP* expression. (c,d) Ear with 1:1 segregation, a single copy insertion outcross. (e,f) An ear with 3:1 segregation, from a heterozygote self-pollination, showing brighter homozygous kernels amongst less-bright heterozygous *Ds-GFP* kernels. (g,h) An outcross ear segregating both the *Ds-GFP* and *C1*/*c1* (anthocyanin) markers.

EarVision.v1 was built with Tensorflow 1 (Abadi *et al*., 2015), a now deprecated version of Google’s deep learning framework. The original pipeline performed best with separately trained models for the different cameras used for imaging in different years, required breaking a full ear image into parts during inference, used a remote server cluster to run, and did not provide the positional information required for the assessment of spatial effects. EarVision.v2 reimplements a similar kernel detection system in PyTorch (Ansel *et al*., 2024), addressing each of these issues. EarVision.v2 uses the Faster R-CNN object detection architecture (Ren *et al*., 2017) to locate kernels of two different classes: fluorescent and non-fluorescent (*Ds-GFP* mutant and wild-type, respectively). After hyperparameter tuning (Appendix S2), the model was trained using 409 ear images (328 training, 81 validation) with manual bounding-box annotations. The input to the system is a projected ear image from the scanner system (Figure 3a). The first output from the system is a set of bounding boxes for the kernels, each with an assigned class (Figure 3b). This is sufficient for performing counts of fluorescent and non-fluorescent kernels and calculating transmission ratios (Data S1). EarVision.v2 additionally outputs spatial information for each kernel in the form of XY coordinates (Figure 3c) determined by calculating the centroid of each bounding box (Figure 3d; see Methods). These coordinates are output as an ‘ear map’ file in .xml format for each image and are used for downstream spatial analyses.

**Figure 3:**
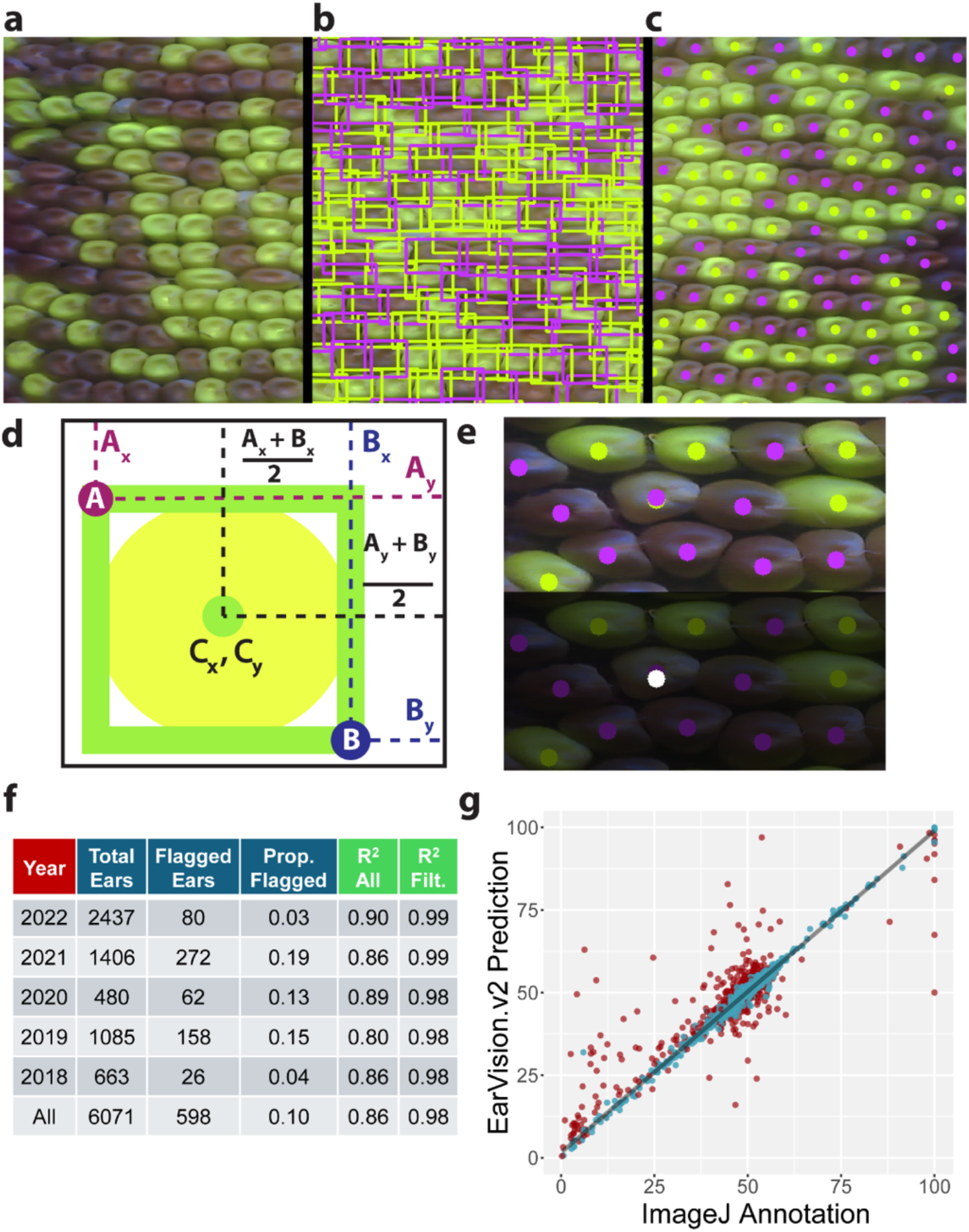
EarVision.v2 computer vision model e4ectively phenotypes and localizes *Ds-GFP* and wildtype kernels. (a) The vision model uses ear projection images as input and (b) outputs bounding boxes for fluorescent and non-fluorescent kernels through an implementation of Faster R-CNN object detection. (c) Centroids of the bounding boxes (green, purple dots) provide a pair of XY coordinates for each kernel, (d) calculated using the top-left [A] and bottom-right [B] corners of the kernel bounding box. (e) Overlapping green and purple centroids (center, top image) define kernels with ambiguous phenotype calls (white dot, bottom image). (f) EarVision.v2 analysis of thousands of ears across multiple field years identifies a minority as ‘Flagged Ears,’, i.e., low quality predictions for subsequent human annotation. R^2^ values from ears with human annotations both with (R^2^ All) and without (R^2^ Filt.) flagged ear images show robust model performance. (g) Transmission rate (%) calculated from human annotations (ImageJ Annotation) plotted against model predictions (EarVision.v2). Blue: Image measurements which pass quality threshold (n=2034). Red: Image measurements which fail quality threshold (n=416).

The calculation of individual kernel coordinates also enabled development of ambiguous kernel detection as an additional quality control feature in EarVision.v2. Ambiguous kernels are detected by determining instances where calculated centroids of the two different classes are within a certain distance from one another (Figure 3e). When such overlap occurs, kernels are marked as ‘ambiguous’, and were further excluded from kernel counts, calculations of transmission, and spatial analysis. A subset of ears had high proportions of such kernels, indicating poor model performance due to a range of factors (e.g., incomplete kernel development, Figure S1a-d). We found that bare patches of unfilled ovules on ears with a small number of kernels were also often difficult for the computer vision model to interpret (Figure S1e,f). By comparing EarVision.v2 predictions to counts from a set of 2450 ears annotated manually using the ImageJ ‘Cell Counter’ tool, we found that ears with fewer than 100 predicted kernels or with more than 2% ambiguous kernels (Figure 3g, red dots; Figure S2; Data S1) were more often associated with inaccurate EarVision.v2 counts. Therefore, we implemented automatic flagging of such ears for manual annotation, recovering accurate data that would not otherwise be usable for analysis. To confirm that model performance metrics were not biased by analysis of training images, we excluded training images with ImageJ annotations and found the overall R^2^ value remained the same (Figure S3). Overall, the EarVision.v2 system achieves transmission rate calculations comparable to human annotation, with an R^2^ value of 0.86 without quality control, improving to 0.98 across all field years when flagged ears are excluded (Figure 3f).

To construct our spatial ear map dataset, we used *Ds-GFP* alleles validated by PCR and Sanger sequencing as insertions into predicted maize gene models (Warman, 2020; Warman *et al*., 2020; Zhou *et al*., 2021), and discarded alleles with less than five independent ear map files for analysis. This dataset includes 58 alleles representing insertions into 54 gene models, encompassing 1384 ear maps (1264 by model inference and 120 by ImageJ annotation; average number of kernels per ear 376, standard deviation 105) and over 500,000 kernels. The majority of ear maps match Mendelian segregation expectations and therefore provide an important control (Data S2). We confirmed that 10 of the 58 alleles are associated with significant pollen transmission defects using a Generalized Linear Model (GLM), which accounts for non-independence of kernels on the same ear (Figure S4, BH-adjusted p≤0.05; Data S3; Warman, 2020; Warman *et al*., 2020; Zhou *et al*., 2021). The GLM estimated transmission rates ranged from 0.10 to 0.46 for the 10 alleles, representing insertions into nine different genes. The associated gene models encode proteins with potential functions in gene regulation, cell wall modification, sperm cell adhesion to target cells, and cell signaling (Data S2). Thus, the alleles appear to span a range of both fitness impacts (subtle to severe) and cellular processes (regulatory to structural) contributing to pollen fertility.

### Spatial inheritance effects are associated with pollen-specific transmission defects

To detect spatial patterns in the analyzed images, we developed a pipeline in R (EarScape) to assess each ear map via statistical modeling (Figure 4). EarScape divides each ear map into 16 bins along each ear, discarding the first and last bin (base and apex) to remove distorted ends of the image projections from consideration. The transmission rate for each bin is calculated from the ratio of fluorescent and nonfluorescent kernels in that bin, providing a spatially ordered set of measurements (Figure 4c). The relationship between the number of fluorescent and nonfluorescent kernels in each bin, and the corresponding X-coordinate of the bin, is then assessed by fitting to two different Generalized Linear Models (GLMs). The first, a ‘Linear’ GLM, fits monotonic patterns which solely increase (base-low, apex-high) or decrease (base-high, apex-low) over the entire ear. The second, a ‘Quadratic’ GLM, fits non-monotonic patterns (such as U-shaped parabolic distributions) but is also able to fit monotonic increasing or decreasing patterns (Figure 5, ear cartoons). The spatial analysis results in three p-values for each ear, helping categorize the type of spatial pattern detected: two p-values derived from the Linear GLM representing the probability of the pattern in the either the increasing or decreasing direction along the ear (apex-high or base-high, respectively, calculated using two opposing one-tailed tests), and one p-value from the Quadratic GLM, calculated using ANOVA (see Methods). The software tool also generates a graph representing each ear map, plotting each bin’s transmission rate and a line representing the fitted GLM (Figure 4d). The pipeline thus provides outputs (quantitative and visual) enabling rapid assessment of particular ears for specific non-random spatial patterns.

**Figure 4:**
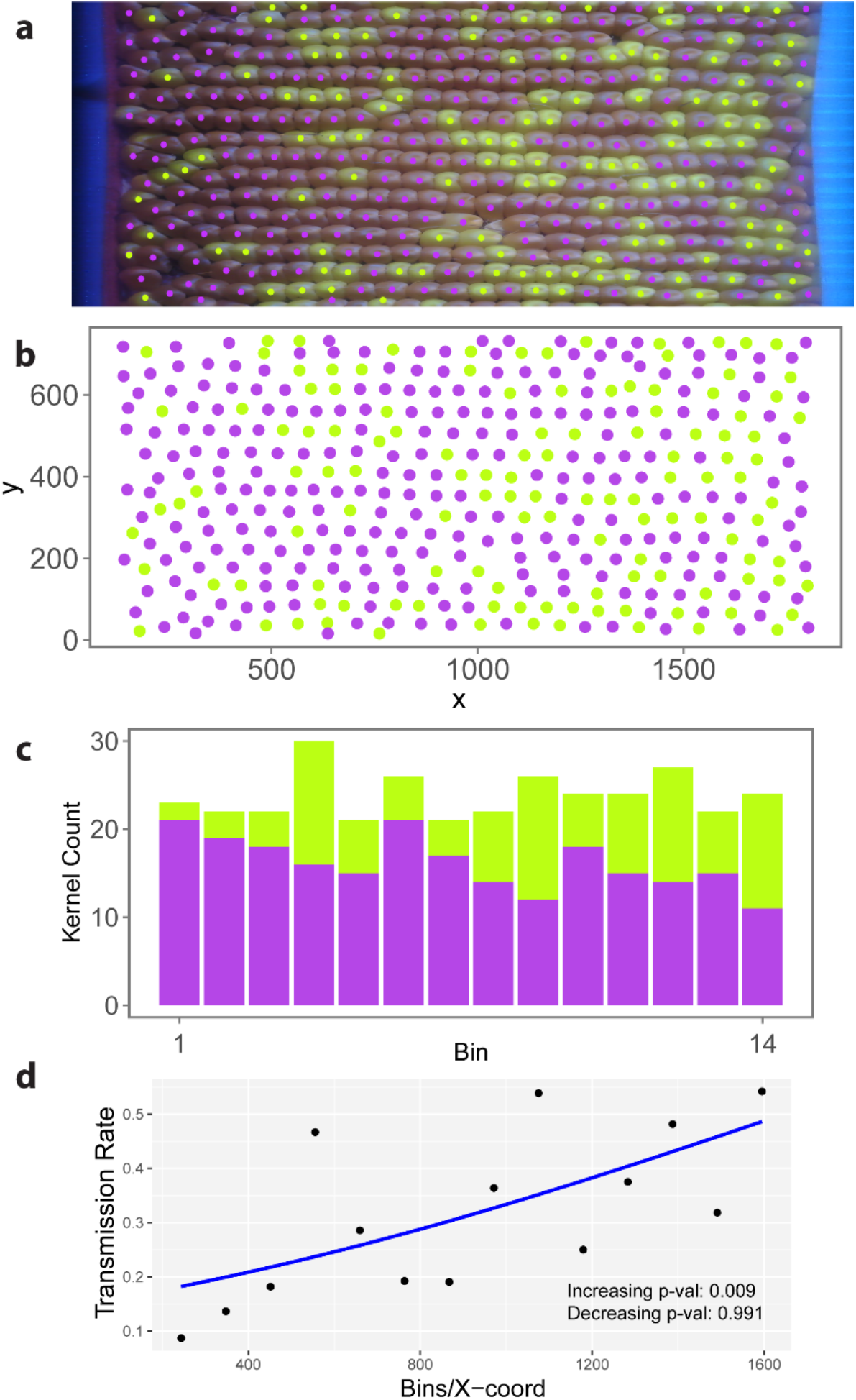
The EarScape statistical pipeline assesses non-random spatial patterns in varying transmission rates across the maize ear. (a, b) The EarVision.v2 vision system provides centroid coordinates for fluorescent and non-fluorescent kernels, oriented base-to-apex left-to-right. Ear is from a pollen cross for the *tdsgR102H01* insertion. Axes: pixel coordinates. (c) The length of the ear is divided into 16 equally-spaced bins (Bin 0 – Bin 15), and the transmission rate is calculated for each bin as Kernel Count GFP (Green) / Kernel Count WT (Purple). Bins 0 and 15 are discarded due to distortion in these regions of the ear projections. (d) Per-bin kernel count data are used to fit ear GLM to detect spatial gradient in transmission rate (blue line). Linear GLM p-value ≤ 0.05 in 1-tailed test shows increasing or decreasing spatial trend diverging from random distribution.

**Figure 5:**
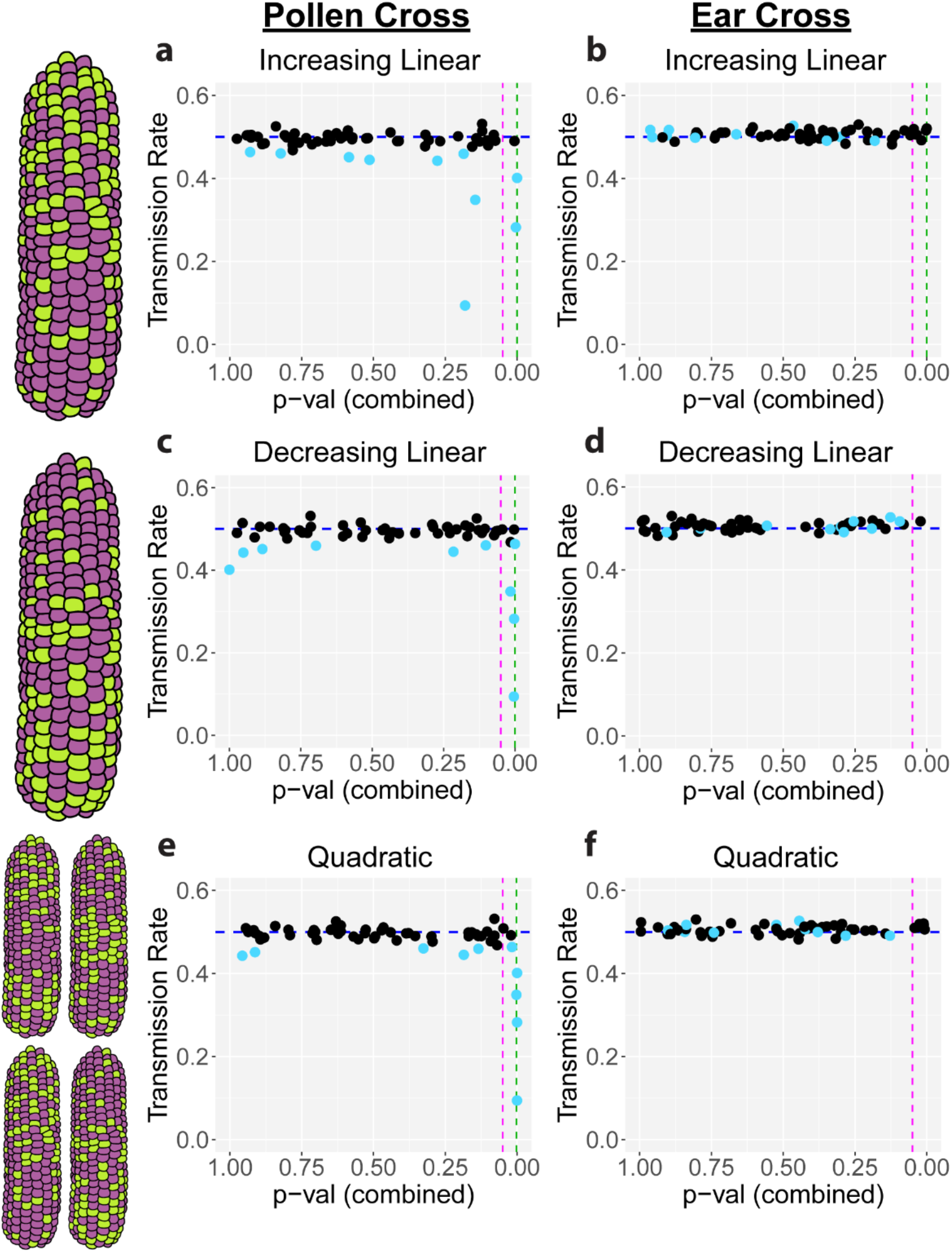
Non-random spatial trends are associated with pollen transmission defects. Plots show p-values for a specific spatial trend vs. estimated transmission rate for each *Ds-GFP* allele, using Fisher’s combination method on GLM-derived p-values for each ear. Black: alleles with Mendelian segregation; Cyan: alleles with significant transmission defects (Fig. S4). (a, c, e) Results from crosses with the *Ds-GFP* insertion transmitted through the pollen, using the *Ds-GFP*/+ parent as the male. (b, d, f) Results from control crosses with *Ds-GFP* insertion transmitted through the ear, using the *Ds-GFP*/+ parent as the female. (a, b) Increasing linear trend (base-low, apex-high) (c, d) Decreasing linear trend (base-high, apex-low). (e, f) Quadratic trend, encompassing a variety of spatial patterns (U-shaped curvilinear distributions, as well as linear patterns in either direction). Green dotted line: Benjamini-Hochberg threshold with FDR of 0.05. Magenta dotted line: p-value threshold of 0.05.

After applying the EarScape pipeline to the ear maps from all 1384 ears in our dataset, we found ears generated by *Ds-GFP* alleles were associated with a range of spatial patterns (Data S4). In aggregate, 11.8% of ears (65/549) from control crosses, in which any *Ds-GFP* was transmitted through the female gametophyte (hereafter ‘ear cross’) had an unadjusted p value ≤0.05 for at least one of the three statistical tests. In comparison, 18.6% of all experimental ears (155/835), in which any *Ds-GFP* allele was transmitted through the pollen (hereafter ‘pollen cross’) had an unadjusted p value ≤0.05. These results provided evidence that our EarScape analysis identifies significant spatial patterns at a rate above the background, and that the high variability of this biological phenomenon may require a large number of observations to characterize trends within individual alleles.

To characterize spatial trends at the level of individual *Ds-GFP* allele, we used Fisher’s combined probability test (see Methods) to analyze the EarScape p-values (Data S5). We tested the hypothesis that a given allele is associated with a spatial trend using the p-values from multiple independent tests, giving an allele-specific combined p-value for each spatial trend. This analysis identified five alleles with reproducible and significant spatial trends when transmitted through pollen (Figure 5a,c,e), with combined p-values below the 0.05 False Discovery Rate controlling for multiple testing. In control ‘ear’ crosses, these same alleles had neither transmission defects nor evidence of spatial trends (Figure. 5b,d,f), showing that the relationship between the transmission defect and spatial trend in these alleles is pollen specific. Spatial trends were associated with severity of the transmission defect, with all four of the most severe alleles showing a spatial effect detected by the Quadratic model (Figure 5e). Surprisingly, we detected one *Ds-GFP* allele (*tdsgR06C04*, an insertion into a gene coding for a SUMO-conjugating enzyme) with a statistically significant increasing linear trend when used as a female parent (Figure 5b).

This allele did not show a significant transmission bias as either a female (52.2% GFP) or male (49.2% GFP) parent and did not show any spatial effect when used as a male parent (Data S3, S4).

Aggregating data from our full panel of ears generated by *Ds-GFP* allele transmission through pollen, and applying a multiple testing threshold, we detected non-random spatial trends exclusively in alleles with pollen transmission defects (Figure 6). Although *Ds-GFP* alleles with Mendelian transmission make up 83% (48/58) of our panel, we did not find any statistically significant evidence of a spatial pattern associated with insertion alleles in these lines. Of the 17.24% (10/58) of alleles with reduced transmission included in our panel: 10% (1/10) had a significant increasing linear trend, 10% (1/10) had a significant decreasing linear trend, 40% (4/10) had a significant quadratic trend. Overall, 50% (5/10) of alleles with reduced transmission, representing mutations in four of nine genes, can generate spatial trends in progeny seeds on the ear. These results indicate that, unlike alleles inherited in a Mendelian manner, alleles affecting pollen fitness can often contribute to non-uniform spatial patterns of fertilization across the maize inflorescence.

**Figure 6:**
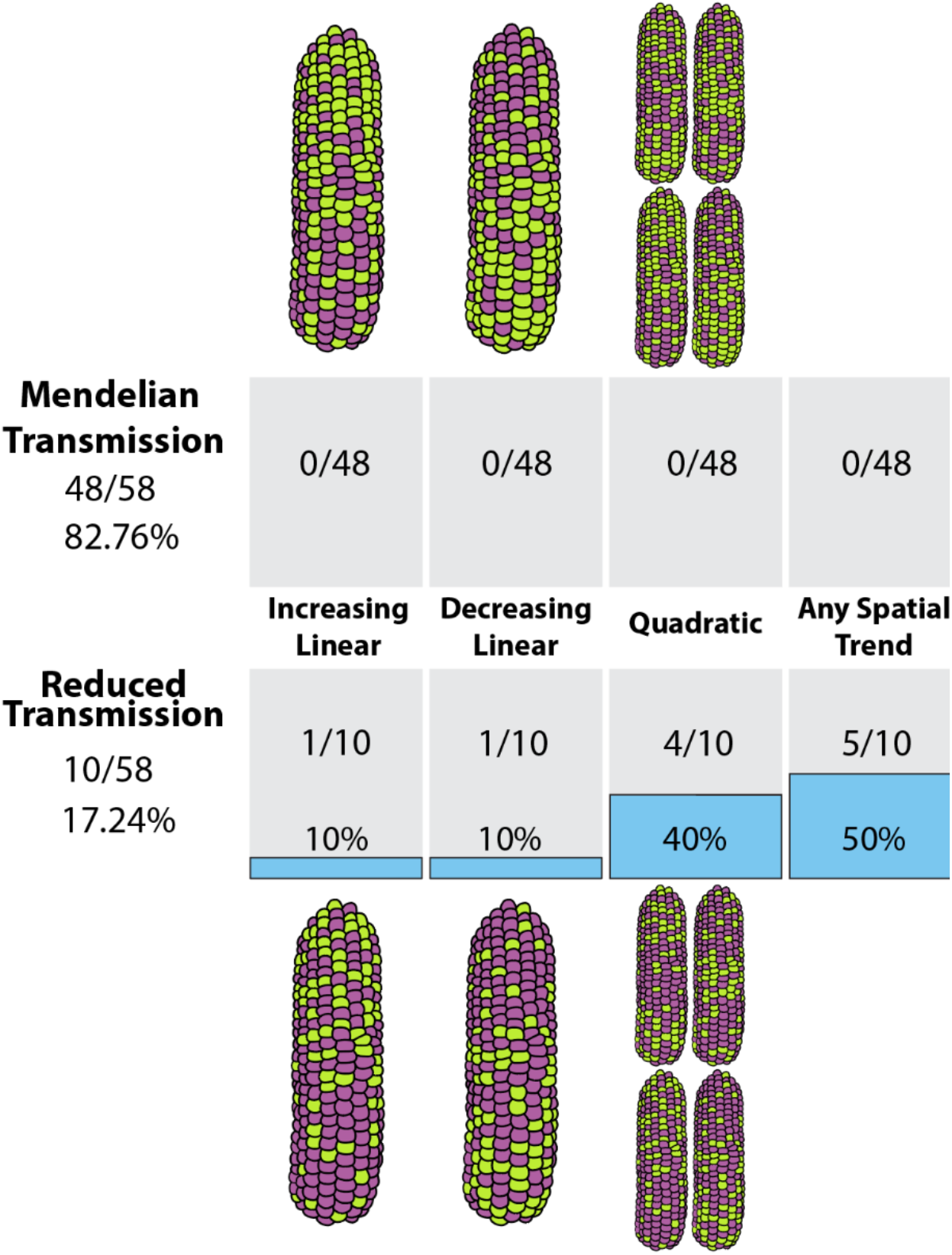
Only alleles with reduced transmission show statistically significant spatial trends when used as a pollen parent. Top: No alleles exhibiting Mendelian transmission had significant spatial trend model fits. Bottom: Five alleles out of ten with reduced male transmission had significant spatial trends. (Two alleles with significant spatial eQects are mutations in the same gene, *gex2*). Data shown from pollen crosses only (where *Ds-GFP*/+ is used as male parent). Counts shown are alleles with p-values passing the Benjamini-Hochberg threshold with FDR = 0.05.

### Statistical modeling indicates that individual alleles can be associated with specific spatial trends

A closer evaluation of the statistical analyses for all ten *Ds-GFP* alleles with pollen transmission defects reveals that the spatial modeling approach can distinguish different categories of the dominant spatial pattern conditioned by each allele (Figure 7). Overall, we detected strong overlap between alleles identified by quadratic trend modeling and those identified by at least one type of linear trend modeling, suggesting that quadratic line fit can capture a range of increasing, decreasing, or curvilinear patterns on specific ears, as each of these patterns can be represented by a different range of a quadratic curve. For example, *Ds-GFP* allele *tdsgR84A12*, which is an insertion into Zm0001eb099800, also known as the sperm cell-specific gene *gex2* (Warman *et al*., 2020), displayed statistical evidence for both increasing linear and decreasing linear spatial trends (combined p-values ≤ 0.01), captured as a significant result for the quadratic trend. On the other hand, *tsdgR102H01*, a *Ds-GFP* insertion in Zm0001eb283600, displayed significant results for increasing linear and quadratic trends only. These results show that although quadratic trend modeling captures multiple pattern types, certain alleles primarily condition a specific spatial pattern, revealed by evaluation of the Linear model in opposing directions. Thus, *tdsgR102H01* and *tdsgR96C12* can be categorized as conditioning ‘increasing’ or ‘decreasing’ patterns, respectively, whereas the other three alleles influence spatial distributions with less consistent patterns. Notably, the five alleles with significant spatial effects alter gene models predicted to function in a variety of cellular roles: RNA binding/regulation (*larp6c1*), sperm cell function at double fertilization (*gex2*), actin cytoskeleton-related (Zm00001eb283600, encoding a putative actin-binding protein, now designated *bag1\**, see below), and cell wall-related (Zm00001eb236740, encoding a putative arabinogalactan protein).

**Figure 7:**
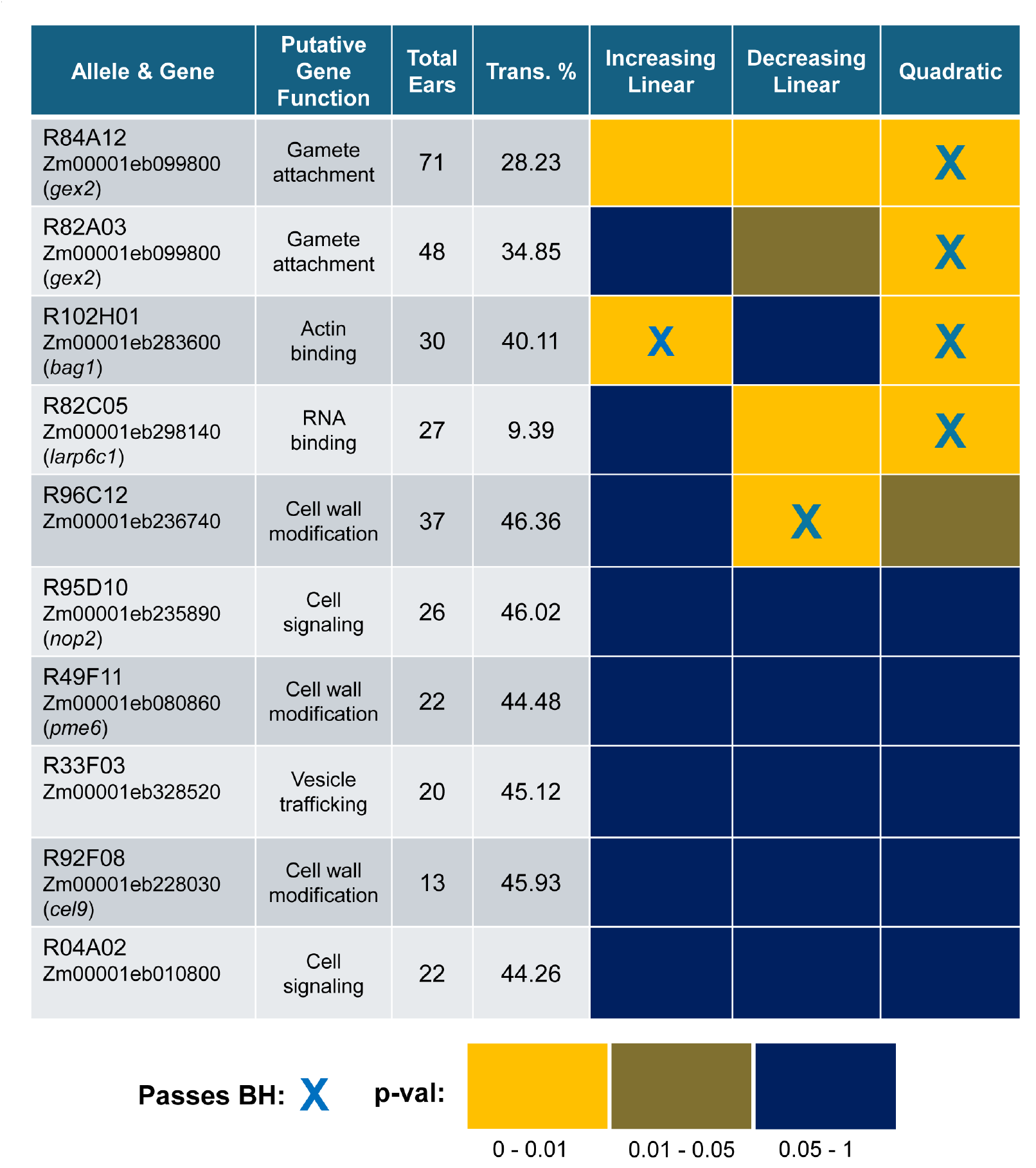
Statistical tests for ten alleles with transmission defects indicate that specific alleles can be categorized with particular spatial trends, across a variety of gene functions. P-value heatmap represents evidence for each evaluated trend for each allele. X: Allele passes Benjamini-Hochberg test with FDR = 0.05. Orange: Unadjusted Fisher combined p-value in range 0 – 0.01. Brown: Unadjusted Fisher combined p-value in range 0.01 – 0.05. Blue: Unadjusted Fisher combined p-value in range 0.05 – 1.

### Spatial inheritance trends vary on a per pollination basis

To better understand the connection between spatial trends at the allele level and patterns observed in each individual pollination, we performed an ear-by-ear post-hoc analysis, assigning each ear to the exclusive categories of ‘increasing linear’, ‘decreasing linear’, or ‘true quadratic’ (see Methods; Figure 8; Data S4, S6), where ‘true quadratic’ ears are those with spatial trends that are only significant under the Quadratic model, and typically are associated with a non-monotonic, parabolic U shape in either direction (Figure 8c,e). The ‘true quadratic’ pattern is rare and only detected in ∼2% of ears analyzed (22/835 pollen crosses, 11/549 ear crosses; Data S4, S6), further supporting the idea that although quadratic modeling is effective at detecting a wider range of spatial effect types, linear modeling captures the majority of spatial effects with greater specificity.

**Figure 8:**
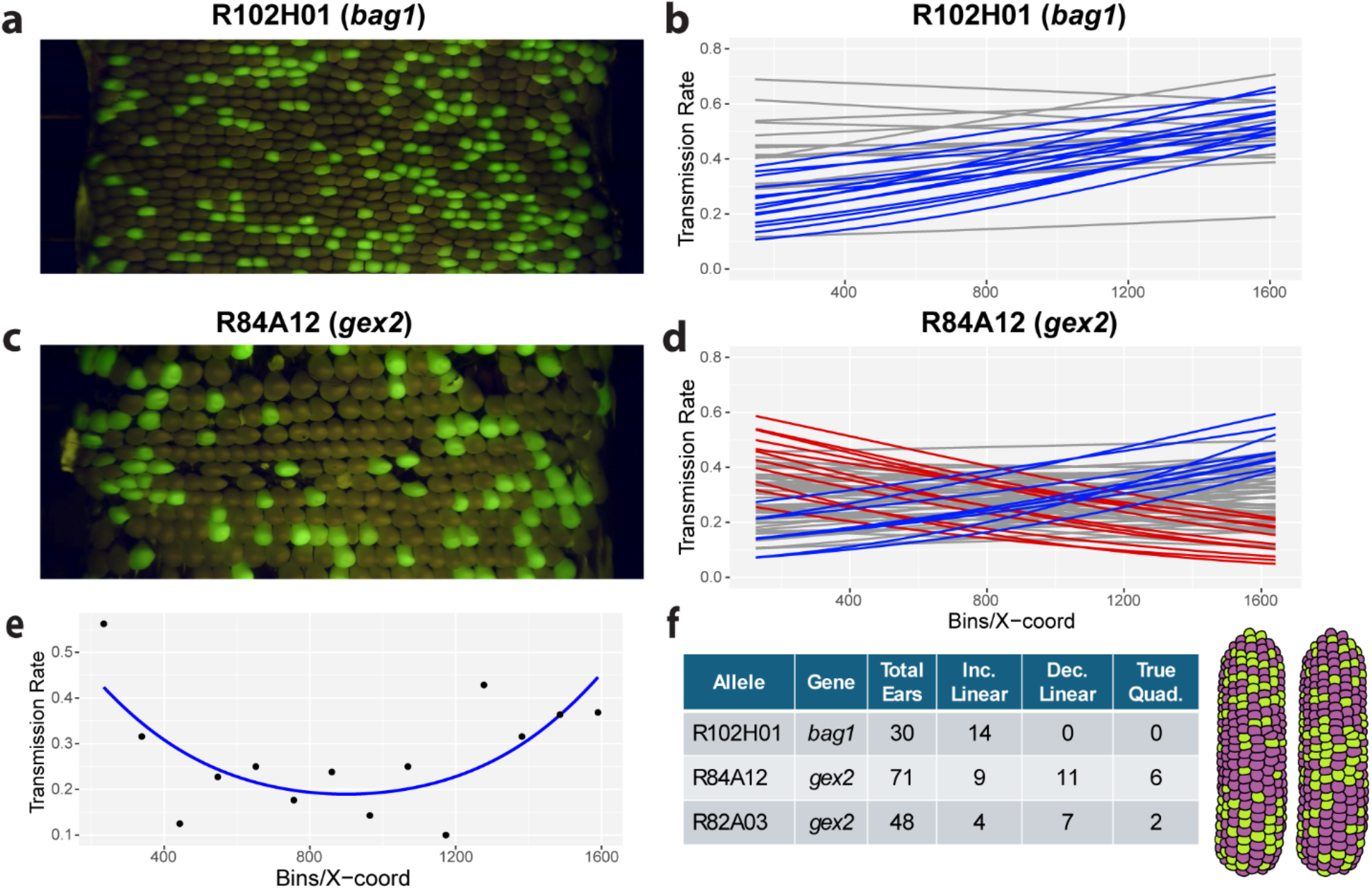
Although particular spatial trends are associated with specific alleles, ears generated from the same allele can show heterogeneous spatial patterns. (a) Ear from pollen cross for the allele *tdsgR102H01*, an insertion in *bag1*, showing increased fluorescent kernels from base to apex. (b) Superimposed Linear GLM pattern lines for all 30 pollen crosses with *tdsgR102H01* show approximately half of the ears are associated with a significant increasing pattern (i.e., increased transmission at the ear apex). Blue: p-value ≤ 0.05 increasing linear test, Grey: not significant. (c) Ear from pollen cross for allele *tdsgR84A12*, an insertion in *gex2*, showing increased fluorescent kernels on both ends of ear. (d) Superimposed Linear GLM pattern lines for all 72 pollen crosses with *tdsgR84A12* show significant increasing and decreasing patterns on separate ears. Blue: p-value ≤ 0.05 increasing linear test, Red: p-value ≤ 0.05 decreasing linear test, Grey: not significant. (e) Quadratic GLM fit for *tdsgR84A12* ear image shown in (c), an example of a ‘true quadratic’ ear (f) Summary of spatial pattern heterogeneity for three *Ds-GFP* insertion alleles. Cartoon shows ‘true quadratic’ ears.

Examining inheritance of *Ds-GFP* across individual ears revealed that the alleles with transmission defects consist of a mix of ears with and without significant spatial patterns (Figure S5), including both increasing and decreasing patterns. Allele *tdsgR102H01* differed from all others analyzed, in that, when associated with a significant spatial effect in a particular ear, it conditioned effects that only fit the linear increasing model (Figure 8a,b). Moreover, the frequency of ears showing significant spatial effects was highest for this allele: nearly half of ears assessed (14 of 30 total ears; Figure 8f) had increasing linear post-hoc spatial patterns. Due to the consistency of this effect, we have preliminarily designated the *tdsgR102H01*-associated gene *base-to-apex gradient1\** (*bag1\**), as no second allele is yet available to confirm this gene as causal for the phenotype. Because of its increasing linear trend, this phenotype suggests *bag1*\* impacts pollen tube competition in the longest silks growing from the base of the ear.

In contrast, the other four alleles with significant spatial effects yielded individual ears that were placed into both increasing and decreasing categories (Figure S5), and in all cases the frequency of ears with spatial pattern was lower than with *bag1*-tdsgR102H01*. Of particular note are the two independent alleles of *gex2, tdsgR84A12* and *tdsgR82A03*, which both show a similar overall mixture of ear types, despite their quantitative differences in severity (*tdsgR84A12* transmits at 28%, whereas *tdsgR82A03* transmits at 35%). Both alleles produced ears with significant increasing linear, decreasing linear, and true quadratic spatial patterns (Figure 8c,d,e,f). The frequency of the ‘true quadratic’ type of pattern was notably higher for these alleles than in the overall dataset (8.5% and 4.2% for *tdsgR84A12* and *tdsgR82A03*, respectively), indicating that loss of *gex2* function in the sperm cells is particularly likely to condition this unusual pattern. Overall, although the likelihood of a spatial pattern appears to be driven primarily by altered pollen fitness, the range of spatial outcomes observed do not consistently reflect the expectations of pollen competition shaped by genotype-driven differences in pollen tube growth.

## Discussion

In 1926, Mangelsdorf and Jones reported biased inheritance of the *sugary1* marker depending on kernel position, with differential inheritance in the apical vs. basal halves of the ear (Mangelsdorf and Jones, 1926; Emerson, 1934). They hypothesized that this spatial pattern might be explained by alleles at the linked *Gametophytic factor1* (*Ga1*) locus, differentially influencing pollen tube growth rate based on gametophytic genotype. We now know that alleles of *Ga1* can create a reproductive barrier by inhibiting pollen tube growth in incompatible silks and, in some cases, promote apex-biased seed set (House and Nelson, 1958; Lausser *et al*., 2010; Lu *et al*., 2014; Lu *et al*., 2020).

Despite this early observation, few genetic loci have been identified in maize which spatially bias inheritance patterns based on pollen genotype (Gorla and Rovida, 1980; Pollak *et al*., 1995; Fowler, 2003). Although spatial effects have been reported in several other plant species, including *C. pepo* (Quesada et al., 1991) and *Arabidopsis* (Meinke, 1982), these reports are complicated by multiple genetic differences between cultivars or only characterize the effects of variants at a single gene. A prevailing hypothesis in these studies is that slower pollen tube growth rates lead to a competitive disadvantage as the distance from the stigma to the embryo sac increases.

However, spatial bias in fertilization success could arise from a range of mechanisms beyond pollen tube growth rate. For example, differential pollen tube attrition in the style (Barnes and Cleveland, 1963; Lausser *et al*., 2010; Lu *et al*., 2014), or spatial heterogeneities across floral structures, such as variation in ovule maturity or embryo sac development, may interact with pollen genotype to determine fertilization success. To address these questions more broadly, we created a high-throughput spatial ear imaging and analysis platform and applied it to a panel of *Ds-GFP* insertion alleles.

### EarVision.v2 with EarScape identifies spatial pollination effects in high throughput

The updated version of our phenotyping platform, EarVision.v2, improves the design of the imager and generalizes the computer vision model to increase throughput and scale of the available dataset. Critically, EarVision.v2 also outputs the relative location of each categorized kernel on the ear, which we used for statistical analysis of spatial effects. Determining kernel location also enabled a feature to computationally identify ambiguous kernel classifications. This not only improves accuracy for each image but also allows automatic identification of ear images that are recalcitrant to computer vision assessment (i.e., those with >2% ambiguous kernels). Such images can either be annotated manually, or removed from the overall dataset, as appropriate based on ear quality. The effort invested into platform improvement enabled generation of a suitably large set of ears with phenotyped and ‘mapped’ kernels for statistical analysis via the EarScape pipeline.

In previous studies, testing for a spatial effect on inheritance relied on comparing transmission rates at two contrasting regions, for example progeny at the apex vs. base of the *Arabidopsis* silique (Meinke, 1982; Cole *et al*., 2005). EarScape quantifies allele transmission in multiple regions and then models transmission distributions across the entire ear. Using both linear and quadratic models increases the potential for EarScape to detect subtle or unexpected patterns. The variation observed in our dataset indicates that a large number of quantitative observations and rigorous multiple testing correction are needed to confidently detect spatial fertilization patterns.

### Determinants of pollen function interact with individual ears to generate spatial bias

Our analysis found that spatial bias in inheritance is exclusively associated with five of ten *Ds-GFP* alleles associated with pollen fitness defects. Moreover, we found that the four alleles with the most severe transmission defects all also condition spatial effects, suggesting that the more severe the impact on pollen function, the more likely a genotype will result in a spatial bias on inheritance across the ear. Thus, our results support the idea that genetic defects that reduce pollen fitness often generate significant spatial patterns in the distribution of progeny from a heterozygous parent.

We detected considerable heterogeneity in spatial patterns across the ears analyzed. Even the allele most consistently associated with a spatial effect, *bag1*-tdsgR102H01*, yielded a statistically significant spatial bias in about half of the ears measured. We speculate that this variability results from interactions with the environment, such as differences in floral structure maturity at the time of pollination, or from technical constraints, such as worker-to-worker differences in pollination technique. Our high throughput analysis of inheritance allowed us the statistical power to detect allele-conditioned spatial effects despite ear-to-ear variability.

Given the prevailing hypothesis that differences in pollen tube growth rate drive post-pollination male gametophyte competition (Brink and MacGillivray, 1924; Correns, 1928; Walsh and Charlesworth, 1992; Williams and Reese, 2019), we expected “base-low, apex-high,” transmission bias, designated here as an increasing pattern, would be the most abundant spatial bias in our dataset. However, only *bag1*-tdsgR102H01* was best characterized by an increasing trend consistent with the pollen tube growth hypothesis. This is particularly surprising in maize, given its extraordinarily long style and the many competing pollen tubes per ovule (Heslop-Harrison *et al*., 1985). We found the opposite, “base-high, apex-low,” decreasing pattern was apparent in ears generated by pollination with each of the other four alleles associated with spatial biases, and the predominant outcome for *tdsgR96C12*. Post hoc analysis determined that the two *gex2* alleles (*tdsgR84A12* and *tdsgR82A03*) produced the highest percentage of ears with the ‘true quadratic’ curvilinear pattern (Fig. 8e). Thus, although certain alleles generate predominant spatial effects (*bag1*-tdsgR102H01*, increasing), other alleles yield patterns that depend on the individual pollination. Thus, the distribution of kernels on each ear can be seen as an interaction between pollen genotype and the specific ear being pollinated.

### Possible mechanistic links between pollen alleles and spatial pattern

Our current data allow speculation about mechanisms underlying the observed spatial patterns of inheritance. With a ‘increasing’ spatial bias, *bag1*-tdsgR102H01*best fits the original model of decreased pollen tube growth rate limiting competition with wild-type pollen as silk length increases. In support of this hypothesis, all *bag1*-tdsgR102H01*/+ pollinated ears transmit at near-Mendelian rates towards the ear apex (Fig. 8b), suggesting that the earliest period of pollen tube growth is not significantly different from that of wild-type. Given that the actin cytoskeleton plays a critical role in pollen tube tip growth (Cole and Fowler, 2006; Fu, 2010; Zhang *et al*., 2023), and that the candidate *bag1\** gene model (Zm00001eb283600) encodes a predicted actin-binding protein, an attractive hypothesis is that BAG1* provides a cytoskeletal function which facilitates pollen tube growth within the silk.

After our post hoc analysis, we determined that the curvilinear ‘true quadratic’ pattern was most frequently observed in two independent alleles of the *gex2* sperm cell-specific gene. Both alleles can also condition increasing and decreasing patterns (Fig. 8f). GEX2 is thought to encode an extracellular factor promoting adhesion of the sperm cells to their targets in the embryo sac, the egg and central cells (Mori *et al*., 2014). The range of patterns seen from the *gex2*/+ pollinations raises the possibility that sensitivity of the egg and central cells to GEX2-defective sperm is dynamic, perhaps associated with a particular maturation stage of the ovule or embryo sac. In this view, the specific *gex2* inheritance pattern on an ear would reflect ovule maturity across the ear at the time of pollination. Testing each of these mechanistic hypotheses will require specific experiments assessing each – e.g., manipulating silk length to either relieve or exacerbate the *bag1*-tdsgR102H01* spatial effect.

## Conclusion

By generating a high-throughput analytical pipeline and statistically characterizing a large number of alleles, our results broaden the types of pollen-conditioned spatial patterns of inheritance documented in flowering plants. Given previous work, we anticipated that the base-low, apex-high trend would be the most common, based on the idea that pollen tube growth rate is a major determinant of spatial bias in inheritance. However, our data suggest *in vivo* pollinations integrate complex interactions between pollen gametophytic genotype and the carpel to produce increasing, decreasing, and curvilinear patterns.

At least in one case, these findings contradict assumptions derived from *in vitro* pollen tube growth experiments. The shorter pollen tubes observed *in vitro* for *larp6c1-tdsgR82C05* mutants (Zhou et al., 2021) predict an increasing spatial pattern. However, *in vivo* spatial effects for *larp6c1-tdsgR82C05* were best fit by the Quadratic model, with individual ears exhibiting decreasing as well as increasing patterns. We propose that spatial heterogeneities in addition to silk length, not replicated *in vitro*, can influence final outcomes on the ear. Hence, our approach, identifying subtle yet reproducible spatial phenotypes associated with pollen mutants, should help inform our understanding of how specific gametophytic genes function *in vivo*.

We suggest that it may be useful to view the broader than expected range of spatial inheritance patterns documented here from an ecological perspective (Heslop-Harrison, 2000; Harder *et al*., 2016). Consider thousands of genetically distinct seeds distributed across a landscape and allowed to grow. The eventual success of the individuals would reflect the distribution of genotypes and their innate potential, but also the interaction between genotype and types of shade, soil or geography found in the environment. In a pollination, the recipient flower can be viewed as a landscape with a range of heterogeneities, such as silk length and ovule maturity in maize, that interact with pollen to favor particular genotypes, generating spatial patterns after fertilization. As more mutant alleles are functionally characterized, commonalities and mechanisms underlying their spatial effects should also help identify the specific factors that govern gamete competition within this floral landscape.

## Materials and Methods

### Ds-GFP Insertion Lines Cultivation and Pollination

*Ds-GFP* alleles were acquired and cultivated at the Oregon State University Botany and Plant Pathology Farm in Corvallis, OR as in Warman et al 2020. Briefly, lines were requested from the Maize Genetics Cooperation Stock Center in which genes expressed highly in pollen, based on RNA-seq data, harbored insertions into the gene model, based on data from acdsinsertions.org (Xiong et al., 2013; Data S2). All insertion alleles in this study had been previously validated via PCR followed by Sanger sequencing to confirm insertion coordinates (Warman, 2020; Warman *et al*., 2020; Zhou *et al*., 2021). For generating experimental crosses with specific *Ds-GFP* alleles, GFP-positive kernels from outcrosses to the W22-*r c1* inbred line (i.e., heterozygous plants) were grown to maturity, sampled for DNA, and genotyped using allele-specific primers paired with primers internal to the *Ds-GFP* element (Data S7). These GFP-positive heterozygotes were used as parents for controlled reciprocal pollinations to either to the W22-*r c1* inbred line, or to an alternate a *c1* homozygous, predominantly W22 tester line according to standard practices (Neuffer, 1994). Briefly, fresh pollen was insured by setting tassel bags on the day of pollination in the morning following removal of already exserted anthers, and were taken down one to two hours later, with all collected pollen used to saturate the silks on recipient ears. To increase pollen competition, silks on recipient ears were allowed to grow back for two to three days after cutting, generating extensive silk surface area for capturing applied pollen. After each harvest, all ears were dried and organized for imaging before shelling for long term storage.

### Scanner Platform and Imaging

The components to make MES.v2 can be inexpensively purchased from common online retailers, and the system can be assembled following the diagram, 3D printer files, and parts list included in Supporting Materials (Appendix S1). Briefly, the design and construction was accomplished by undergraduate capstone project Team 616 in the Oregon State College of Engineering (MIME ME497-ME498), with the goal of improving throughput from the original MES (Warman et al 2021) without a wholesale redesign (e.g., the same rotating ear approach, general lighting and digital imaging parameters, and ear scanning workflows were used as in the original). The digital camera was updated to an ELP 5-50mm Varifocal Lens 1080P USB Camera with H.264 High Definition Sony IMX323 Webcam. The code for scanning ears, generating projections, and uploading those into cloud storage was also updated and is available at https://github.com/fowler-lab-osu/EarScannerUtilities.

The set of ear projections used for the training set included 409 examples from the summer field seasons of 2018, 2019 and 2022, encompassing images generated from three different digital cameras and two different versions of the MES. For this training set, projections were manually annotated using either Labellmg or Label Studio (https://labelstud.io/), with kernels outlined with bounding boxes and labelled as either fluorescent, non-fluorescent or ambiguous. When training the object detection model, kernels annotated as ambiguous are treated as having both a fluorescent and non-fluorescent label. Each image took approximately 10-20 minutes to fully annotate with bounding boxes. To generate the larger set (2450 total) of ear counts that were used to validate EarVision.v2 accuracy, the Cell Counter plug-in in ImageJ/Fiji (https://imagej.net/software/fiji/downloads) was used to tag kernels with the fluorescent, non-fluorescent or ambiguous labels, with the resulting count and spatial location data saved as an .xml format file.

### Model and Model Training

EarVision.v2 (https://github.com/fowler-lab-osu/EarVision2) was trained using the fasterrcnn_resnet50_fpn_v2 Faster R-CNN network (Ren *et al*., 2017) included in the torchvision.models Pytorch subpackage (Torch Contributors, 2022; torchvision.models.detection.fasterrcnn_resnet50_fpn_v2). We used the following values for hyperparameters specific to this implementation: Region proposals retained for training and inference = 3000, region proposals sampled = 512, IOU threshold = 0.7, trainable backbone layers = 4, non-maximum suppression threshold = 0.5, box score threshold = 0.2. A hyperparameter search (Appendix S2) was performed to determine the optimal values for the number of trainable backbone layers, the non-maximum suppression threshold, and box score threshold.

The training dataset included 409 images from the 2018, 2019 and 2022 field years and was segregated into an 80/20% training/validation split (328 training, 81 validation). The network was trained for 30 epochs with a learning rate of 0.005, and the best epoch (27) was chosen based on F1 score and difference in calculated transmission ratio of validation set. Training and inference were performed on a computer workstation with an Nvidia RTX A6000, 48 GB VRAM GPU (Appendix S2).

To estimate transmission rates and identify non-Mendelian alleles, a GLM model to analyze transmission data for the 58 alleles of interest (as generated by EarVision.v2 or manual Fiji annotation) was implemented as in Warman *et al*., (2020). Multiple testing correction for the transmission rate GLM was performed using R’s p.adjust() function with the Benjamini-Hochberg method (method = “BH”).

### Determining Kernel Centroid Coordinates and Ambiguous Kernel Calls

Using the top-left and bottom-right coordinates for the bounding box, the centroid coordinates for a detected kernel is calculated as follows (Figure 3b):

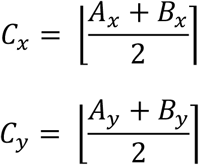

where *C*_*x*_, *C*_*y*_ indicate the centroid coordinates, *A*_*x*_, *A*_*y*_ and *B*_*x*_, *B*_*y*_ the coordinates of the top-left and bottom-right corners of the bounding box, and where ⌊ ⌉ bracket notation indicates half-to-even rounding (Banker’s rounding).

The Faster R-CNN network as implemented in the torchvision.models PyTorch subpackage can output overlapping bounding boxes of different classes if an object plausibly belongs to more than one class. Therefore, in cases where the kernel fluorescence is ambiguous, the system will output boxes for both the fluorescent and non-fluorescent classes. We detect ambiguous kernel calls by finding cases where a small region surrounding the centroid coordinates of two different classes overlap.

For counts of kernel types used for calculating transmission rates, ambiguous kernels are counted by detecting overlapping centroid regions and calculating the area of overlap. Circles with radii of 8 pixels are centered on the centroids for each kernel class, and overlapping regions are detected using a logical conjunction operation (bitwise AND) over two images containing circles marking fluorescent and non-fluorescent kernels. A detected overlapping region with an area greater than 20 pixels is counted as an ambiguous kernel. The centroids of these overlapping regions are also calculated and used in the following step.

For labeling specific ambiguous kernel coordinates in the XY coordinate outputs used for spatial analysis (enabling ambiguous kernels to be ignored in the analysis), each labeled kernel is compared against the ambiguous kernel centroids detected in the prior step. The Euclidean distance between each labeled kernel centroid and each ambiguous kernel centroid is taken, and any labeled kernel with a distance of less than 16 from an ambiguous kernel is considered to be ambiguous. In this case, the ambiguous kernel centroid is added to the output used for spatial analysis (with a label of ‘ambiguous’) if it has not already been included. This ensures that neither of the two original labeled kernels (one fluorescent, one non-fluorescent) which caused the ambiguous kernel call is included, but rather the single ambiguous centroid representing both.

### Determining Spatial Patterns via EarScape

The coordinate data for each ear was divided into 16 equal bins, split vertically. The 1^st^ and 16^th^ bin were discarded due to image distortion at the ends of ears, leaving 14 bins as part of the spatial analysis pattern modeling. Empirical testing of bin number indicated that 16 bins was a good compromise between having enough degrees of freedom for effective modeling and maintaining sufficient kernel numbers in each bin; results did not differ appreciably when using similar numbers of bins. Fluorescent and non-fluorescent kernel counts from bins were used to calculate the relationship between bins and transmission ratio via two models (Linear and Quadratic), using the Generalized Linear Model (GLM) approach to fit spatial patterns using R’s glm() function. The models were of the quasibinomial family and used logit link functions. The function fit by the Linear GLM is:

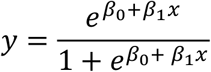

The function fit by the Quadratic GLM is:

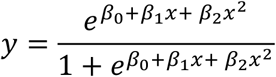

One-tailed tests for assessing increasing and decreasing directionality for the linear pattern were performed using the cumulative distribution function for a t-distribution in R (the pt() function, with n-p-1 (14 – 1 – 1 = 12) degrees of freedom, where n=number of bin observations and p=the number of model parameters) on the t-value returned by the glm summary function for the upper and lower tails respectively. To obtain a p-value representative of the fit for the Quadratic model to an individual ear (admitting both linear and curvilinear patterns), ANOVA was used to compare the Quadratic model and a Null (intercept-only) model. In this framework, the Null model is the reduced model (i.e., fewer terms) and the Quadratic is the full model (i.e., more terms). The ANOVA two-tailed p-value represents the fit of Quadratic model relative to a straight, flat line. The ANOVA compares the test statistic to an F-distribution with 2, 11 degrees of freedom, 2 being the difference in the number of parameters between the Null and Quadratic model and 11 being the degrees of freedom of the full model calculated using n-p-1. EarScape spatial analysis code is available at https://github.com/fowler-lab-osu/EarScape.

### Fisher’s Combination Method

The Linear and Quadratic GLMs return p-values for each ear individually. Pattern GLM p-values for each ear of a given allele were combined using Fisher’s method of p-value combination to obtain a single p-value for each trend for each allele. The test statistic *X*^2^ used to calculate the Fisher’s combined p-value was found by:

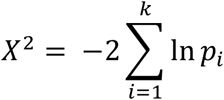

where *p*_*i*_ is the list of p-values. The combined p-value for an allele is obtained by using the R function pchisq() on the test statistic X^2^ with df = 2k… lower.tail=False, log.p=False.

Multiple testing correction for the combined spatial trend p-values was performed by calculating a threshold for significance using the Benjamini-Hochberg procedure with FDR = 0.05 (Benjamini and Hochberg, 1995). The threshold is established at the level of the largest p-value where

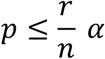

where *r* is the rank of the p-value, *n* is the total number of p-values, and *α* is the FDR. P-values at or below the threshold are considered significant. In instances where no p-value fulfills the criterion for establishing a threshold, no p-values are considered significant.

### Post-Hoc Analysis

For pattern heterogeneity quantification, ears were assigned to categories of ‘increasing linear’, ‘decreasing linear’, and ‘true quadratic’. A ‘true quadratic’ ear is one where the p-value for the Quadratic GLM is ≤ 0.05 and the p-values for both the increasing and decreasing linear patterns is > 0.05, representing a case where a spatial trend is exclusively captured by the Quadratic model (i.e., the Linear model misses it). Ears with an increasing or decreasing linear pattern p-value ≤ 0.05 are assigned to the ‘increasing linear’ or ‘decreasing linear’ categories respectively, regardless of the p-value for the Quadratic model.

## Supporting information

SupportingFigures

Appendix1_MaizeEarScannerv2_DesignFiles

AppendixS2_Computer_Vision_Development_Supplement

DataS1_EarVisionv2_InferenceOutput_5Years

DataS2_58_alleles_kernel_counts

DataS3_transmission_rate_glm

DataS4_earscape_analyses_all_ears

DataS5_allele_results_fisher

DataS6_pattern_counts_per_allele

DataS7_allele_specific_primers

## Acknowledgments

Primary support for the project was to DJ, MM and JF, via NSF grant IOS-2041384. DR is supported by USDA National Institute of Food and Agriculture fellowship 2024-67011-42999. SL is supported by NSF grant IOS-2211434, and USDA National Institute of Food and Agriculture grant 2023-67013-44037. SL, MM and JF are also supported in part by the Oregon Agricultural Experiment Station with funding from the Hatch Act capacity funding program, award numbers NI25HFPORE00216B, from the USDA National Institute of Food and Agriculture. The MES.v2 device was designed and built through a 2020-21 OSU MIME ME497-ME498 capstone project. The students on the project were Tim Pham, Ian Williams, Taylor Amarotico, and Justin Huynh. The course was taught by Prof. Jeff Hoffman and the project was advised by Prof. John Parmigiani. OSU STEM Leaders provided support and orientation for undergraduate researchers LGL and MB. H. Bell, K. Kress, T. Lawrence, J. McDonald, G. Michna, I. Nicholson, A. Perez, C. Waite and H. Woodard provided invaluable assistance for the project in lab and field. Special thanks to H. Dooner and C. Du for generating the *Ds-GFP* insertion population, as well as recommendations and suggestions for effective use of the lines; and for the USDA-supported Maize Genetics Cooperation Stock Center, for distributing seed for *Ds-GFP* stocks.

## Supporting Information

- Supporting Appendices:
  ∘ Appendix S1: MES.v2 Design Files
    ▪ Parts list
    ▪ 3D CAD models/3D printer files
    ▪ Schematics/Diagrams
  ∘ Appendix S2: Computer Vision Development Supplement
- Supporting Data
  ∘ Data S1 – EarVision.v2 Inference output over 5 years (5 tabs)
  ∘ Data S2 – 58 insertion alleles used in spatial analysis: kernel counts for assessing transmission; gene models affected by insertion (2 tabs)
  ∘ Data S3 – Transmission Rate GLM results for 58 alleles (2 tabs, pollen and ear)
  ∘ Data S4 – EarScape analyses, all ears (2 tabs, pollen and ear)
  ∘ Data S5 – Allele results of fisher combined probability test (2 tabs, pollen and ear)
  ∘ Data S6 – Pattern counts per allele (2 tabs, pollen and ear)
  ∘ Data S7 – Gene specific primers for 58 alleles
- Supporting Figures
  ∘ Figure S1 – Examples of images flagged by EarVision.v2’s quality control metrics.
  ∘ Figure S2 – EarVision.v2 performance across 5 years shows that outlier ear projections are effectively flagged by the EarVision.v2 for hand annotation.
  ∘ Figure S3 – EarVision.v2 performance metrics are unaffected when training images are excluded from evaluation.
  ∘ Figure S4 – Analysis of five years of transmission data (>500,000 kernels) for 58 PCR-validated *Ds-GFP* insertions identifies 10 alleles with reduced pollen transmission, at FDR = 0.05.
  ∘ Figure S5 – Superimposed Linear GLM lines, each corresponding to an individual ear, for all 10 transmission defect alleles.
- Code
  ∘ EarVision.v2 Repo: https://github.com/fowler-lab-osu/EarVision2
    ▪ Includes all training set images, with bounding box annotations
    ▪ Includes trained model
  ∘ EarScape Repo (Spatial Analysis): https://github.com/fowler-lab-osu/EarScape
    ▪ Includes coordinate (.xml) files, as well as plots derived from these files, for all ear projection images analyzed
    ▪ Ear projection images (1384 .png files) available upon request
  ∘ EarScannerUtilities Repo: https://github.com/fowler-lab-osu/EarScannerUtilities

## Author Contribution Section

JF, ZV: coordinated field pollinations

JF, ZV, DR: designed data collection procedures

DR: implemented the Faster R-CNN

ML: model training and selection

MB, DJ, DR, LGL, MM: statistics

HF: scanner design and implementation

DR: prepared figures and wrote the manuscript SL, JF, MM: edited the manuscript

