## SupportingFigures for "Spatial inheritance patterns across maize ears are associated with alleles that reduce pollen fitness"

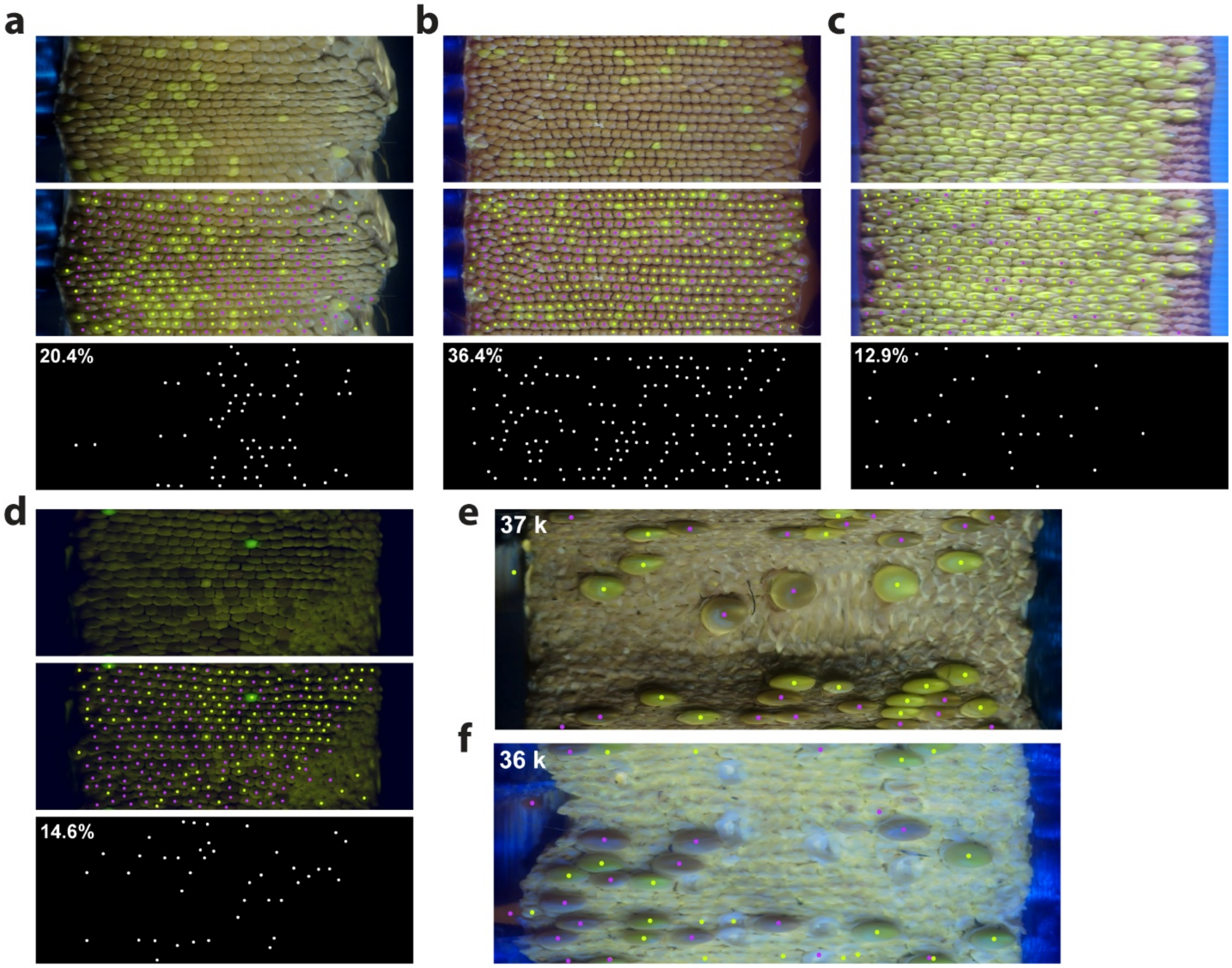


**Supporting Figure 1: Examples of images flagged by EarVision.v2's quality control metrics.** Images with either >2% ambiguous kernels or <100 total non-ambiguous kernels are flagged for hand annotation. (a-d) Ear images flagged due to high percentages of ambiguous kernels. Top: Scanned ear images. Middle: Ear images with annotation centroids. Bottom: White dots where a kernel was called as both fluorescent and non-fluorescent classes, i.e., phenotyped as "ambiguous" by the EarVision.v2. (e,f) Ear images flagged due to low numbers of kernels (k).

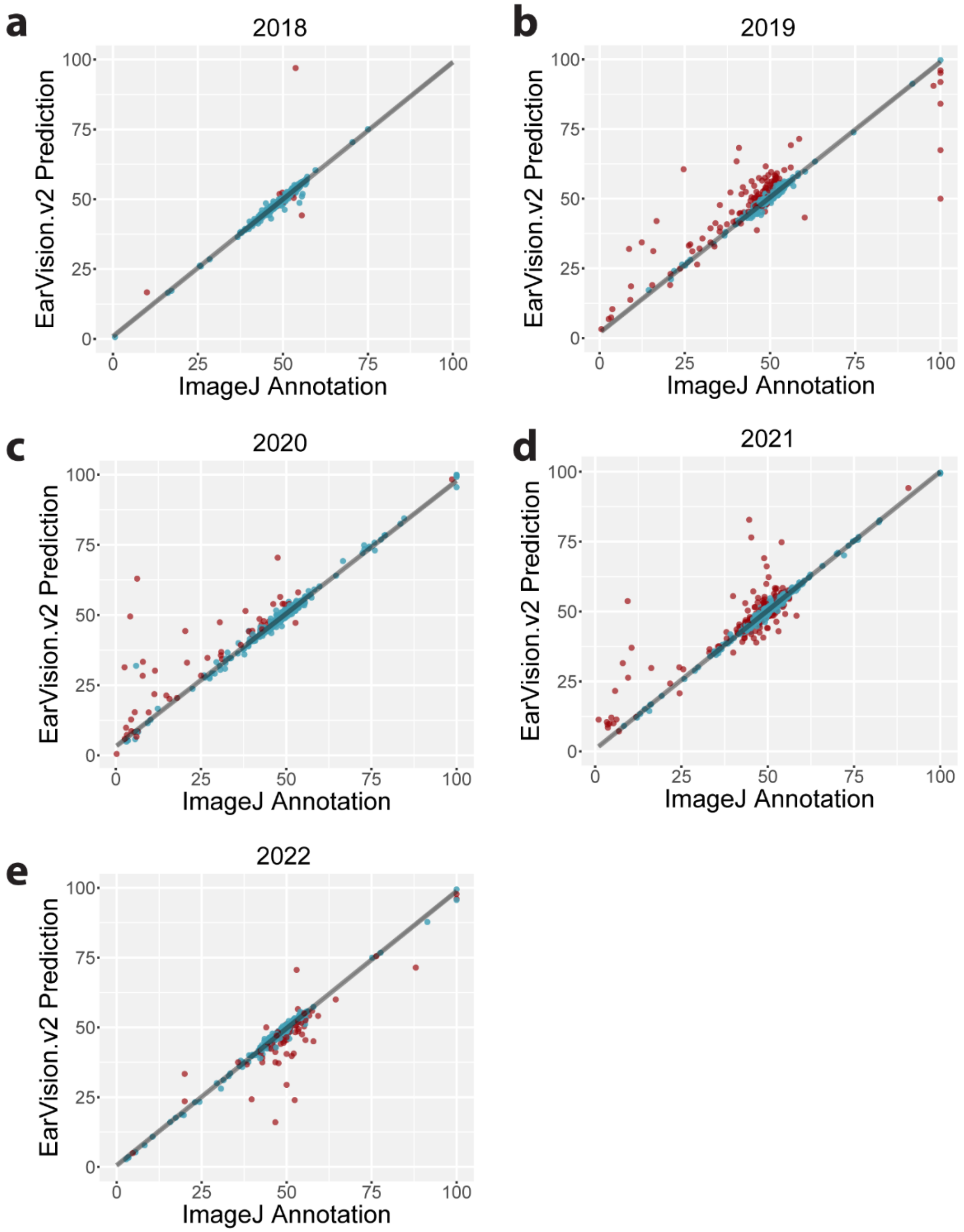


**Supporting Figure 2: EarVision.v2 performance across 5 years shows that outlier ear projections are effectively flagged by the EarVision.v2 for hand annotation.** Transmission rate calculated from human hand annotations ('ImageJ Annotation') plotted against transmission rate calculated from EarVision.v2 kernel predictions. Blue: Image measurements which pass quality threshold. Red: Image measurements which fail quality threshold. Refer to Main Figure 3f for $R^2$ values for each year. (A) 2018 performance data. Blue = 367, Red = 9. (B) 2019 performance data. Blue = 379, Red = 118. (C) 2020 performance data. Blue = 400, Red = 51. (D) 2021 performance data. Blue = 683, Red = 170. (E) 2022 performance data. Blue = 205, Red = 68.

**a**

| Year | Total Ears | Flagged Ears | Prop. Flagged | $R^2$ All | $R^2$ Filt. |
|---|---|---|---|---|---|
| 2022 | 2366 | 72 | 0.03 | 0.90 | 0.99 |
| 2021 | 1406 | 272 | 0.19 | 0.86 | 0.99 |
| 2020 | 480 | 62 | 0.13 | 0.89 | 0.98 |
| 2019 | 977 | 158 | 0.16 | 0.79 | 0.97 |
| 2018 | 560 | 24 | 0.04 | 0.82 | 0.98 |
| All | 5789 | 588 | 0.10 | 0.85 | 0.98 |

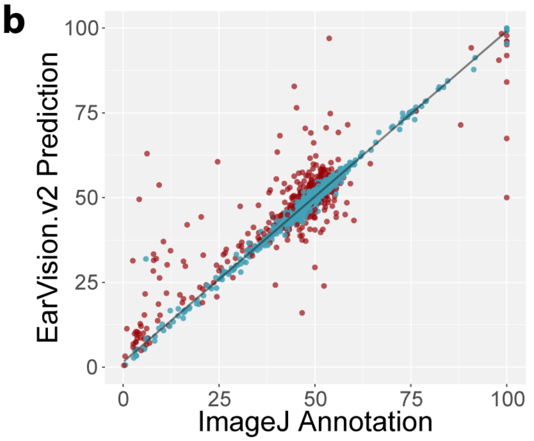


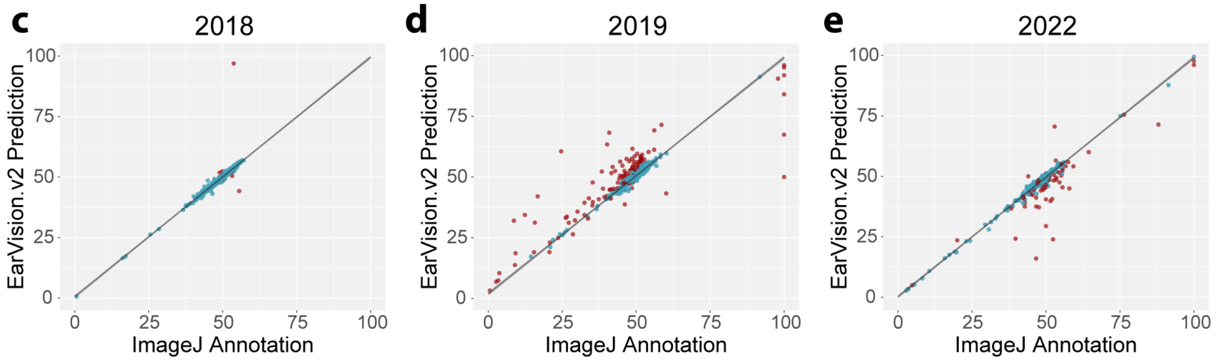


**Supporting Figure 3: EarVision.v2 performance metrics are unaffected when training images are excluded from evaluation.** (a) Removing 282 images with ImageJ annotations that were included in the model training set results in $R^2$ values equivalent to the full set of ears (0.98, Fig. 3f). (b) Transmission rates plotted for five year “held-out” image set (i.e., with training images removed), hand annotation vs. model predictions. Blue: Images passing quality threshold. Red: Images failing quality threshold (c) 2018 performance set with 103 training images removed. (d) 2019 performance set with 108 training images removed. (e) 2022 performance set with 71 training images removed. No images from 2020 and 2021 were included in the training set.

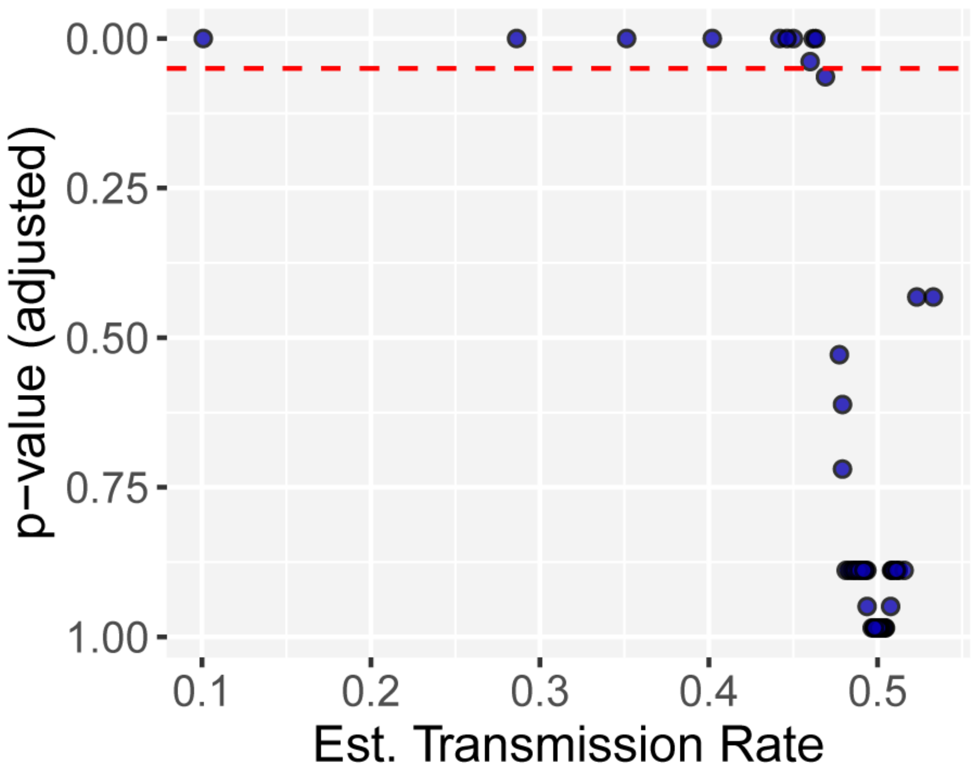


**Supporting Figure 4: Analysis of five years of transmission data (>500,000 kernels) for 58 PCR-validated *Ds-GFP* insertions identifies 10 alleles with reduced pollen transmission, at FDR = 0.05.** Estimated Transmission Rate and multiple-testing corrected p-value calculated from Generalized Linear Model (GLM); red dotted line, Benjamini-Hochberg adjusted p-value ≤ 0.05.

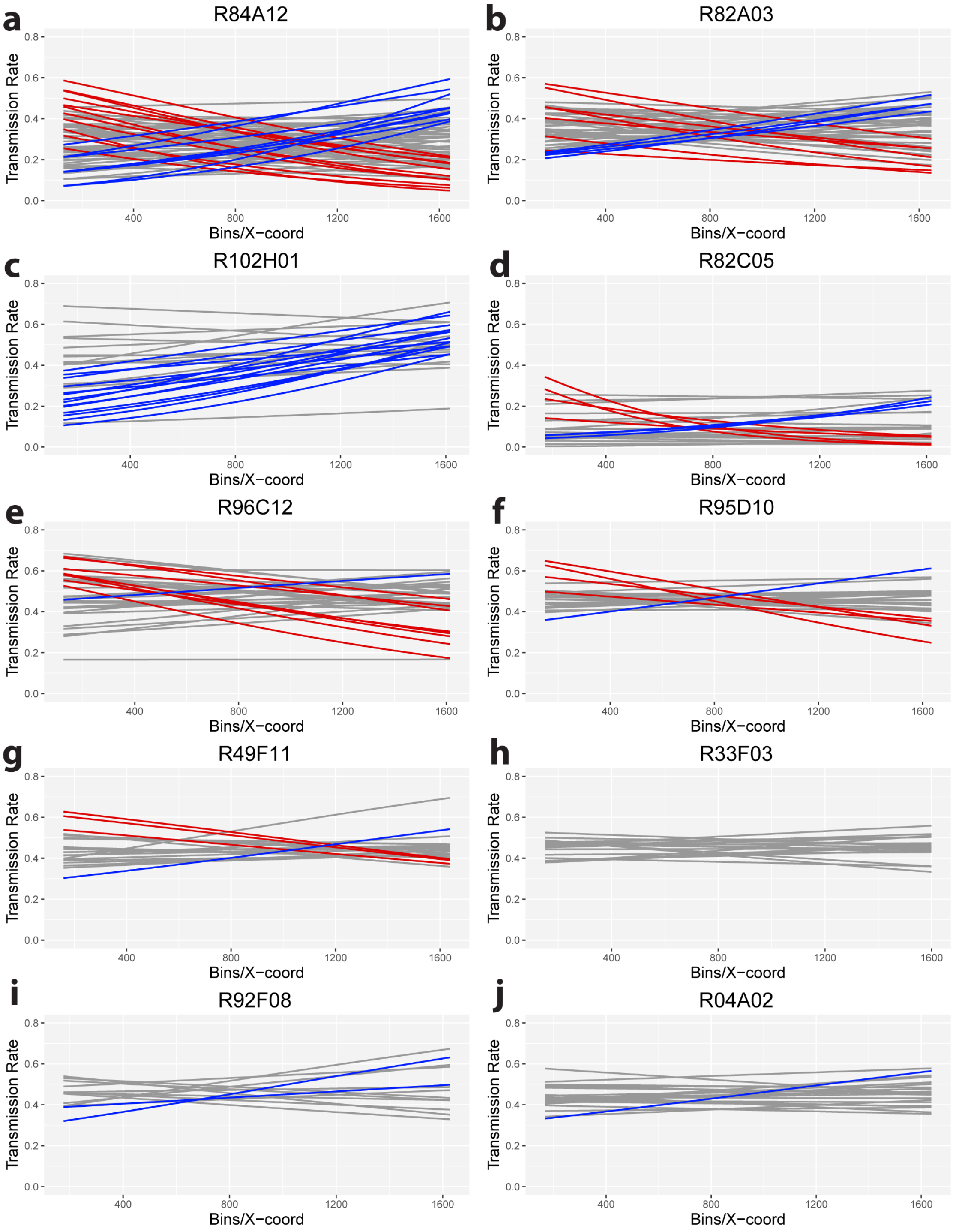


**Supporting Figure 5: Superimposed Linear GLM lines, each corresponding to an individual ear, for all 10 transmission defect alleles.** Blue lines: Linear GLM for ear with p-value ≤ 0.05 for increasing pattern. Red lines: Linear GLM for ear with p-value ≤ 0.05 for decreasing pattern. Grey lines: No significant spatial pattern detected. (A) R84A12 (B) R82A03 (C) R102H01 (D) R82C05 (E) R96C12 (F) R95D10 (G) R49F11 H) R33F03 (I) R92F08 (J) R04A02.
