## Appendix1_MaizeEarScannerv2_DesignFiles for "Spatial inheritance patterns across maize ears are associated with alleles that reduce pollen fitness": MES Assembly Final.pdf

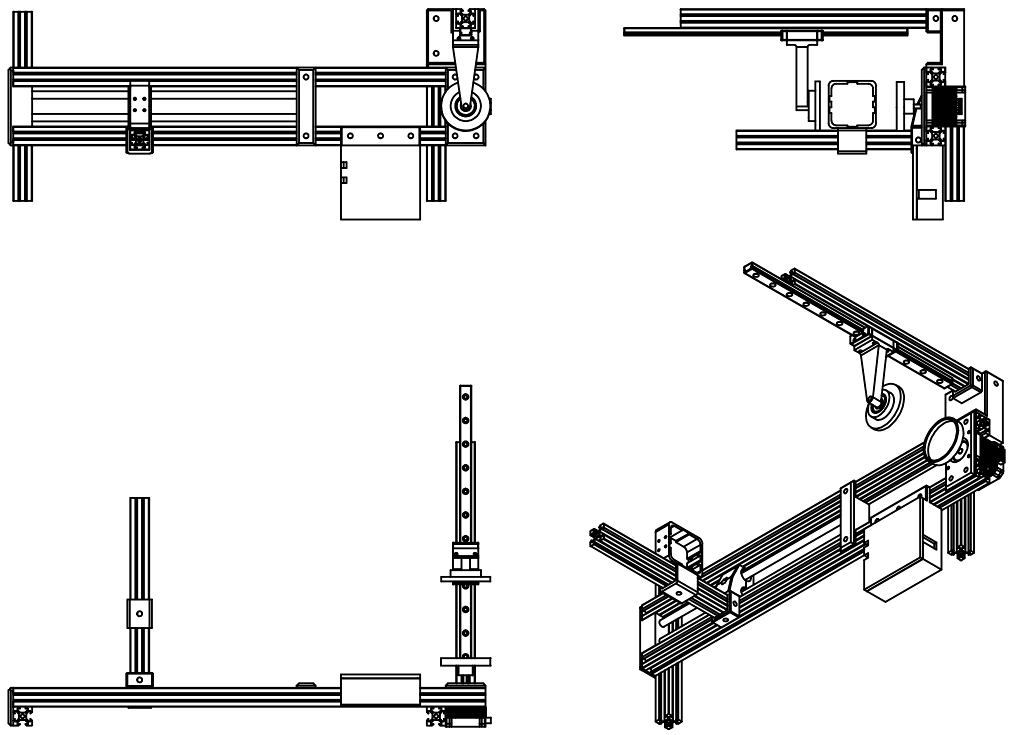

| ITEM NO. | PART NUMBER | QTY. |
|---|---|---|
| 1 | Capstone corn mount adapter | 1 |
| 2 | 608zz | 1 |
| 3 | Camera slide end cap | 1 |
| 4 | Camera slide rear cap | 1 |
| 5 | Camera slider | 1 |
| 6 | Nema17 | 1 |
| 7 | Rotator Top | 1 |
| 8 | Stepper Mount | 1 |
| 9 | 2020-300 | 1 |
| 10 | 2020-500 | 2 |
| 11 | 2020-200 | 3 |
| 12 | mgn12-300 | 1 |
| 13 | 8mm linear rod | 2 |
| 14 | Green End Cap | 2 |
| 15 | Main Block | 1 |
| 16 | Red End Cap | 2 |
| 17 | Main block bearing interface v2 | 1 |
| 18 | MES Bottom foam mount | 1 |
| 19 | MES PCB holder | 1 |
| 20 | Camera_shell_and_slider_v4 | 1 |

DO NOT SCALE DRAWING

| TITLE: MES Assembly Final | | REV 4 |
|---|---|---|
| SCALE: 1:10 | WEIGHT: | SHEET 1 OF 1 |
