## Appendix1_MaizeEarScannerv2_DesignFiles for "Spatial inheritance patterns across maize ears are associated with alleles that reduce pollen fitness": Schematics.pdf

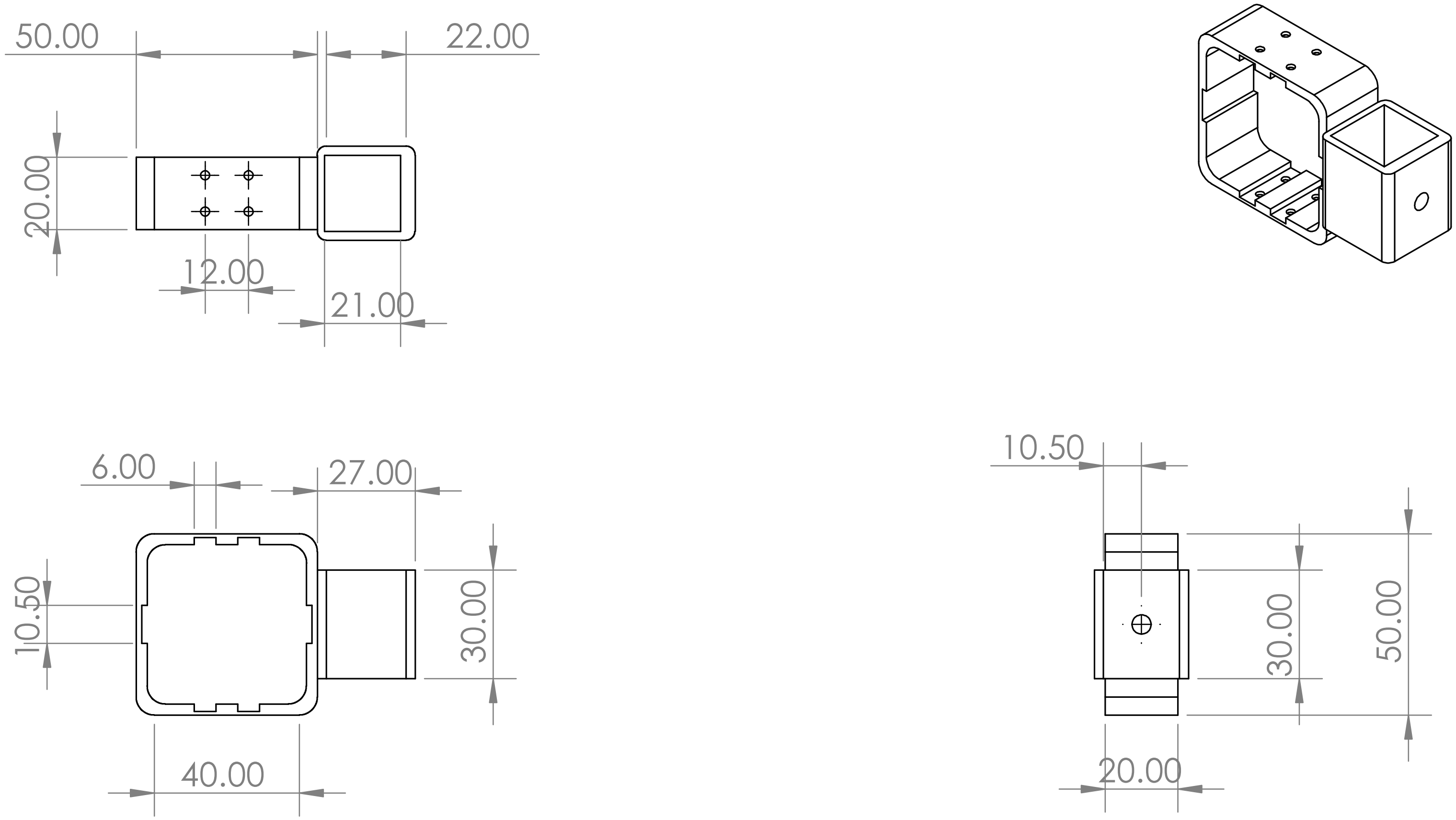

PROPRIETARY AND CONFIDENTIAL

THE INFORMATION CONTAINED IN THIS DRAWING IS THE SOLE PROPERTY OF <INSERT COMPANY NAME HERE>. ANY REPRODUCTION IN PART OR AS A WHOLE WITHOUT THE WRITTEN PERMISSION OF <INSERT COMPANY NAME HERE> IS PROHIBITED.

| NEXT ASSY | USED ON |
|---|---|
| APPLICATION | |

UNLESS OTHERWISE SPECIFIED:

DIMENSIONS ARE IN INCHES
TOLERANCES:
FRACTIONAL ±
ANGULAR: MACH ± BEND ±
TWO PLACE DECIMAL ±
THREE PLACE DECIMAL ±

INTERPRET GEOMETRIC TOLERANCING PER:

MATERIAL

FINISH

DO NOT SCALE DRAWING

| | NAME | DATE |
|---|---|---|
| DRAWN | | |
| CHECKED | | |
| ENG APPR. | | |
| MFG APPR. | | |
| Q.A. | | |

COMMENTS:

TITLE:

Camera_shell_and_slider_v4

SIZE **A** DWG. NO. REV

SCALE: 1:2 WEIGHT: SHEET 1 OF 1

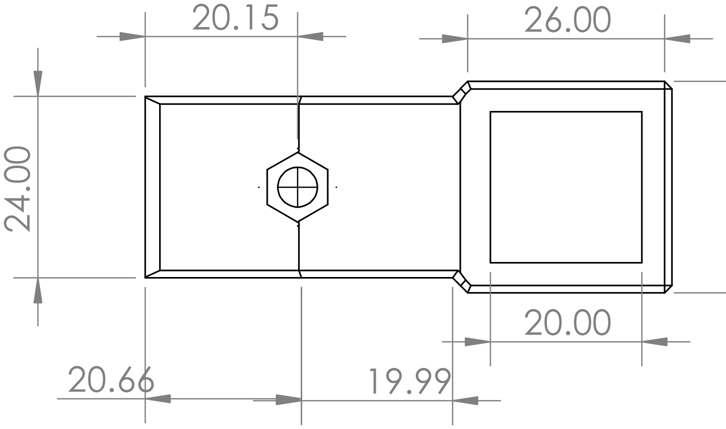

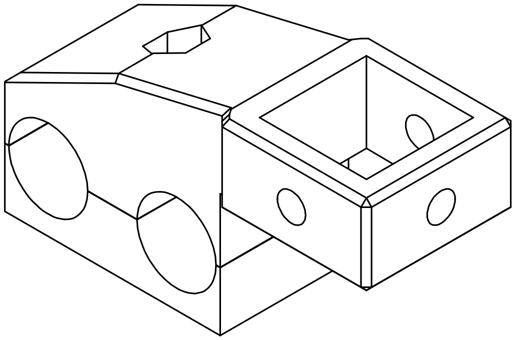

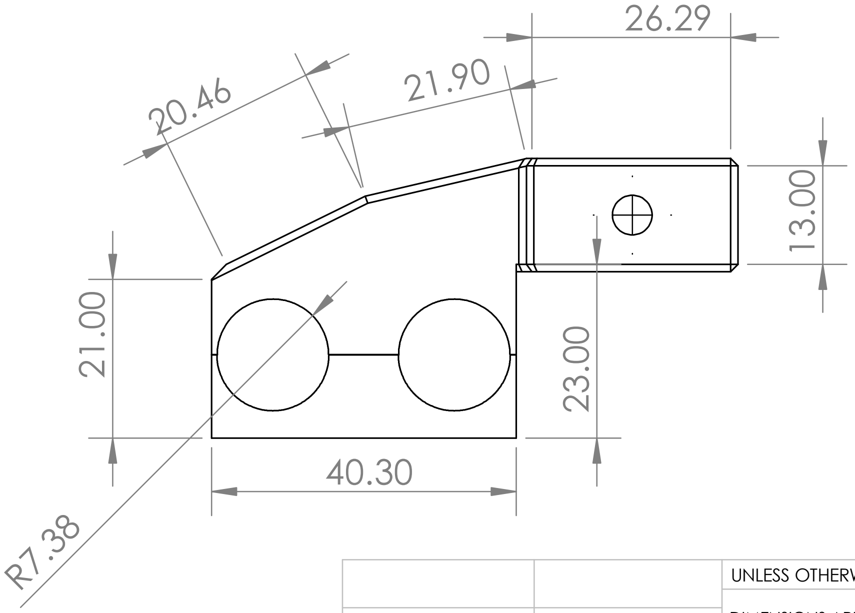

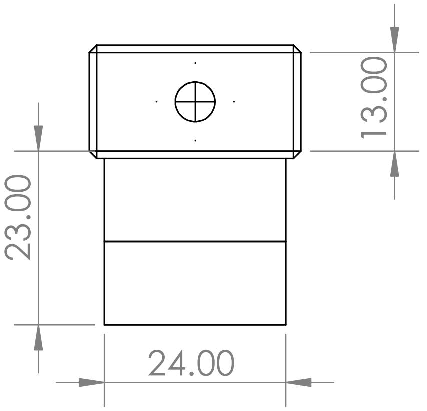

**PROPRIETARY AND CONFIDENTIAL**

THE INFORMATION CONTAINED IN THIS DRAWING IS THE SOLE PROPERTY OF <INSERT COMPANY NAME HERE>. ANY REPRODUCTION IN PART OR AS A WHOLE WITHOUT THE WRITTEN PERMISSION OF <INSERT COMPANY NAME HERE> IS PROHIBITED.

| NEXT ASSY | USED ON |
|---|---|
| APPLICATION | |

UNLESS OTHERWISE SPECIFIED:

DIMENSIONS ARE IN INCHES
TOLERANCES:
FRACTIONAL ±
ANGULAR: MACH ± BEND ±
TWO PLACE DECIMAL ±
THREE PLACE DECIMAL ±

INTERPRET GEOMETRIC TOLERANCING PER:

MATERIAL

FINISH

DO NOT SCALE DRAWING

| | NAME | DATE |
|---|---|---|
| DRAWN | | |
| CHECKED | | |
| ENG APPR. | | |
| MFG APPR. | | |
| Q.A. | | |

COMMENTS:

TITLE:

### Camera slider

| SIZE | DWG. NO. | REV |
|---|---|---|
| **A** | | |

SCALE: 1:1 WEIGHT: SHEET 1 OF 1

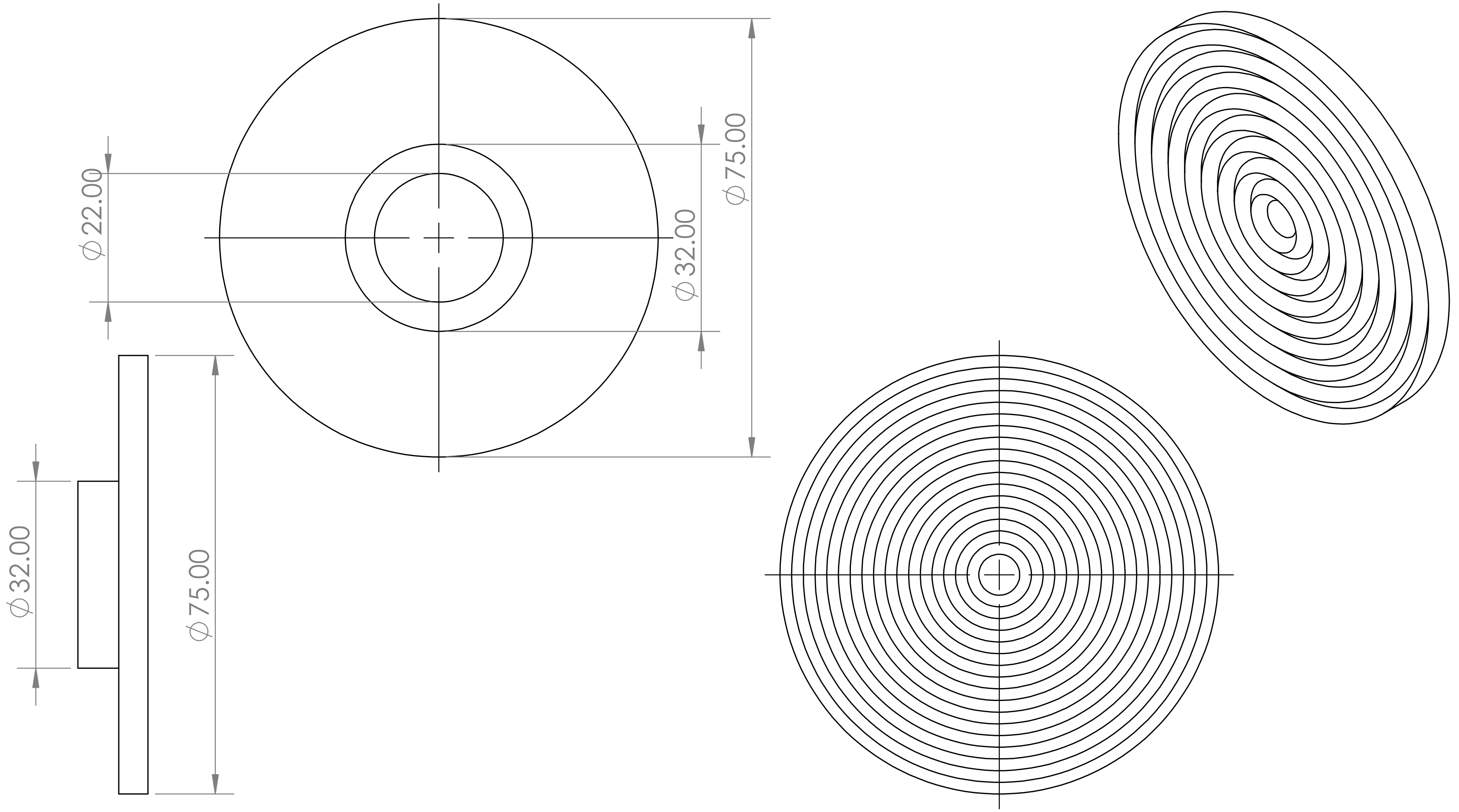

| | |
|---|---|
| NEXT ASSY | USED ON |
| APPLICATION | |

UNLESS OTHERWISE SPECIFIED:

DIMENSIONS ARE IN INCHES
TOLERANCES:
FRACTIONAL ±
ANGULAR: MACH ± BEND ±
TWO PLACE DECIMAL ±
THREE PLACE DECIMAL ±

INTERPRET GEOMETRIC TOLERANCING PER:

MATERIAL

FINISH

DO NOT SCALE DRAWING

| | NAME | DATE |
|---|---|---|
| DRAWN | | |
| CHECKED | | |
| ENG APPR. | | |
| MFG APPR. | | |
| Q.A. | | |

COMMENTS:

TITLE:

### Rotator Top

SIZE **A** | DWG. NO. | REV

SCALE: 1:1 | WEIGHT: | SHEET 1 OF 1

**PROPRIETARY AND CONFIDENTIAL**

THE INFORMATION CONTAINED IN THIS DRAWING IS THE SOLE PROPERTY OF <INSERT COMPANY NAME HERE>. ANY REPRODUCTION IN PART OR AS A WHOLE WITHOUT THE WRITTEN PERMISSION OF <INSERT COMPANY NAME HERE> IS PROHIBITED.

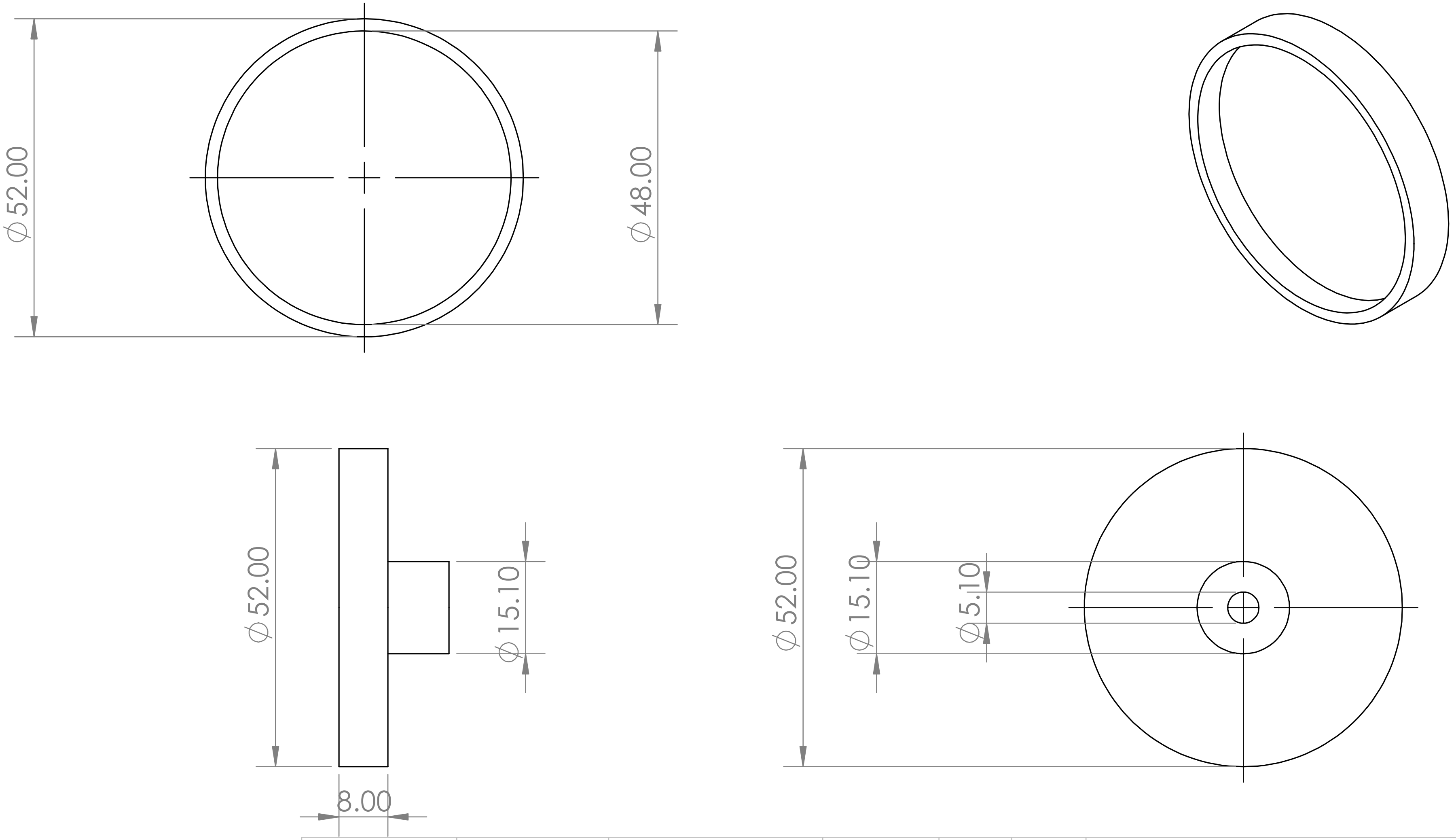

| | |
|---|---|
| NEXT ASSY | USED ON |
| APPLICATION | |

UNLESS OTHERWISE SPECIFIED:

DIMENSIONS ARE IN INCHES
TOLERANCES:
FRACTIONAL ±
ANGULAR: MACH ± BEND ±
TWO PLACE DECIMAL ±
THREE PLACE DECIMAL ±

INTERPRET GEOMETRIC TOLERANCING PER:

MATERIAL

FINISH

DO NOT SCALE DRAWING

| | NAME | DATE |
|---|---|---|
| DRAWN | | |
| CHECKED | | |
| ENG APPR. | | |
| MFG APPR. | | |
| Q.A. | | |

COMMENTS:

TITLE:

MES Bottom foam mount

| SIZE | DWG. NO. | REV |
|---|---|---|
| A | | |

SCALE: 1:1 WEIGHT: SHEET 1 OF 1

**PROPRIETARY AND CONFIDENTIAL**

THE INFORMATION CONTAINED IN THIS DRAWING IS THE SOLE PROPERTY OF <INSERT COMPANY NAME HERE>. ANY REPRODUCTION IN PART OR AS A WHOLE WITHOUT THE WRITTEN PERMISSION OF <INSERT COMPANY NAME HERE> IS PROHIBITED.
