## Appendix1_MaizeEarScannerv2_DesignFiles for "Spatial inheritance patterns across maize ears are associated with alleles that reduce pollen fitness": MES_Basic.ino

```
#define DIR 2
#define STEP 3
#define MS1 7
#define MS2 6
#define EN 8
#define PDN 4
#define BTN1 10
#define BTN2 11

const long duration = 30*1000; // Duration of spin in milliseconds
long stepTime = ((duration*1000) / (200*16)) / 2;

void setup() {
  Serial.begin(115200);
  pinMode(DIR, OUTPUT);
  pinMode(STEP, OUTPUT);
  pinMode(MS1, OUTPUT);
  pinMode(MS2, OUTPUT);
  pinMode(EN, OUTPUT);
  pinMode(PDN, OUTPUT);
  pinMode(BTN1, INPUT_PULLUP);
  pinMode(BTN2, INPUT_PULLUP);

  digitalWrite(EN, HIGH); // Disable stepper on startup. Stepper active LOW
  digitalWrite(PDN, HIGH); // Disable UART
  digitalWrite(MS1, HIGH); // Set microstepping config
  digitalWrite(MS2, HIGH);
}

void loop() {
//  Serial.print(digitalRead(BTN1));
//  Serial.println(digitalRead(BTN2));
  if (!digitalRead(BTN1)) rotate(true);
  if (!digitalRead(BTN2)) rotate(false);
  delay(50);
}

void rotate(bool clockwise){
  digitalWrite(EN, LOW); // Enable stepper motor
  digitalWrite(DIR, clockwise);
  delay(50);
  for (int i = 0; i <= 200*16; i++){
    digitalWrite(STEP, HIGH);
    delayMicroseconds(stepTime);
    digitalWrite(STEP, LOW);
    delayMicroseconds(stepTime);
  }
  digitalWrite(EN, HIGH); // Enable stepper motor
}
```
