## Appendix1_MaizeEarScannerv2_DesignFiles for "Spatial inheritance patterns across maize ears are associated with alleles that reduce pollen fitness": MES_Summary.pptx

### Slide 1
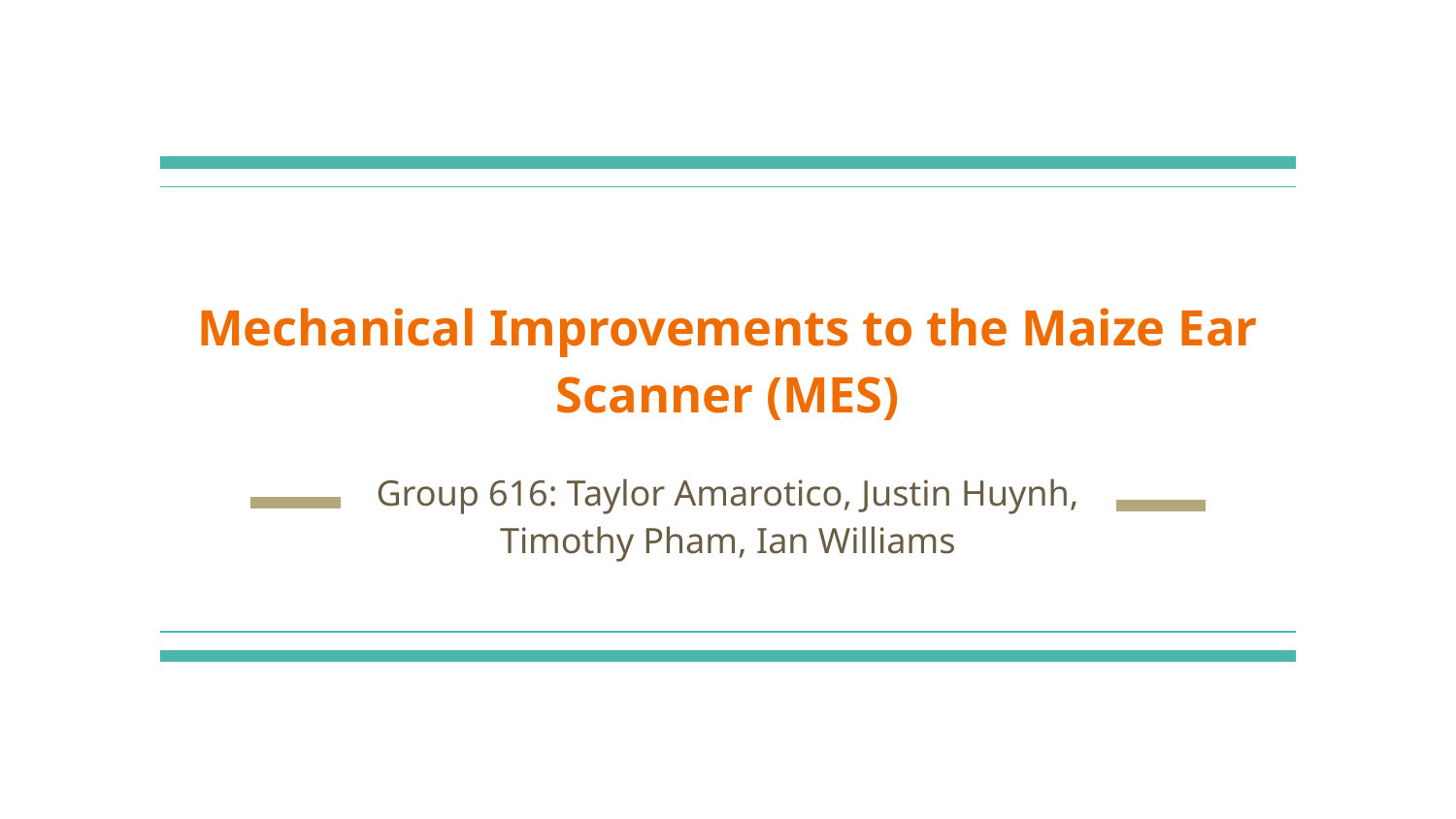

#### Mechanical Improvements to the Maize Ear Scanner (MES)
Group 616: Taylor Amarotico, Justin Huynh, Timothy Pham, Ian Williams

### Slide 2
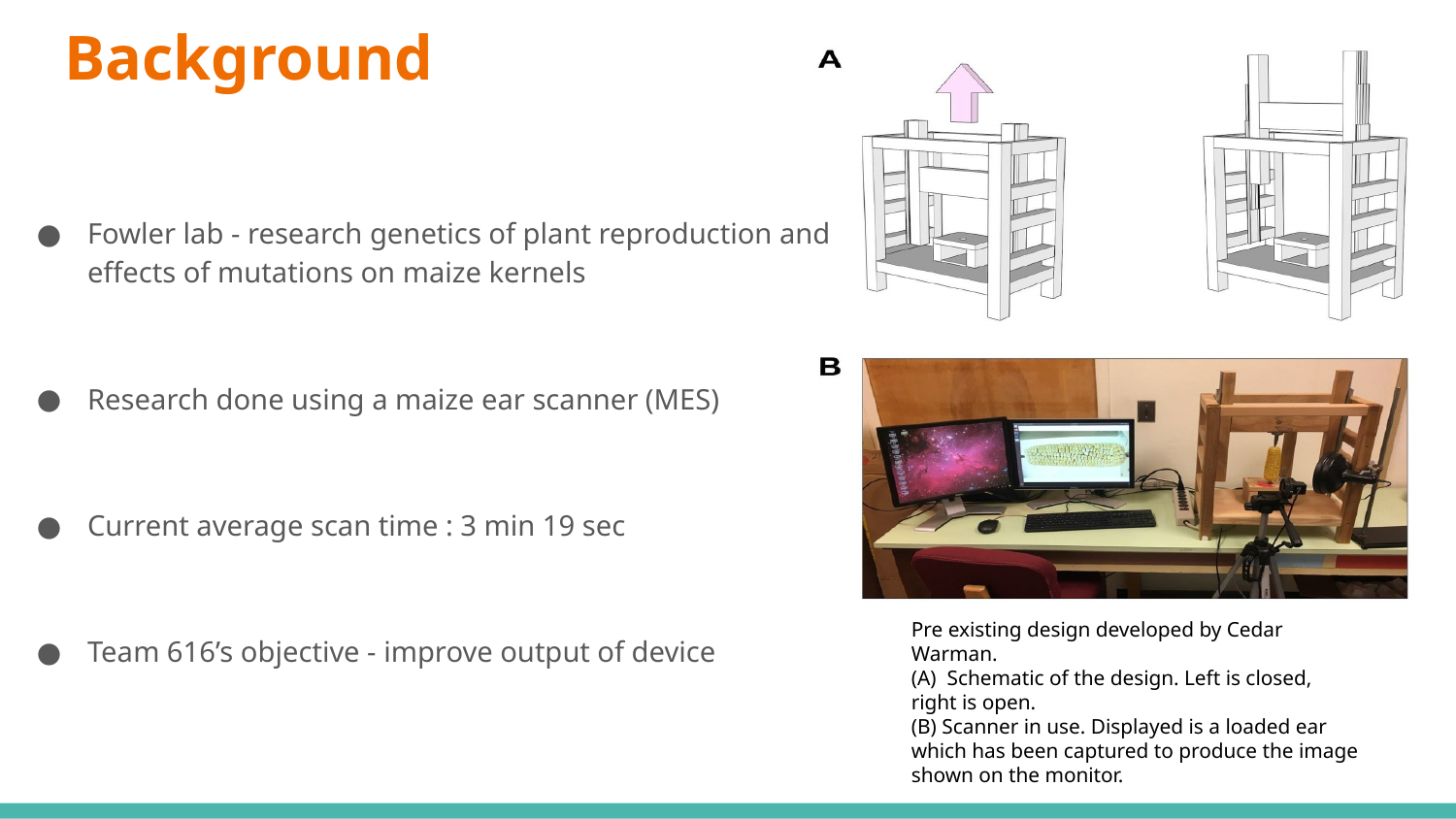

#### Background
Fowler lab - research genetics of plant reproduction and effects of mutations on maize kernels
Research done using a maize ear scanner (MES)
Current average scan time : 3 min 19 sec
Team 616’s objective - improve output of device
Pre existing design developed by Cedar Warman.
(A) Schematic of the design. Left is closed, right is open.
(B) Scanner in use. Displayed is a loaded ear which has been captured to produce the image shown on the monitor.

### Slide 3
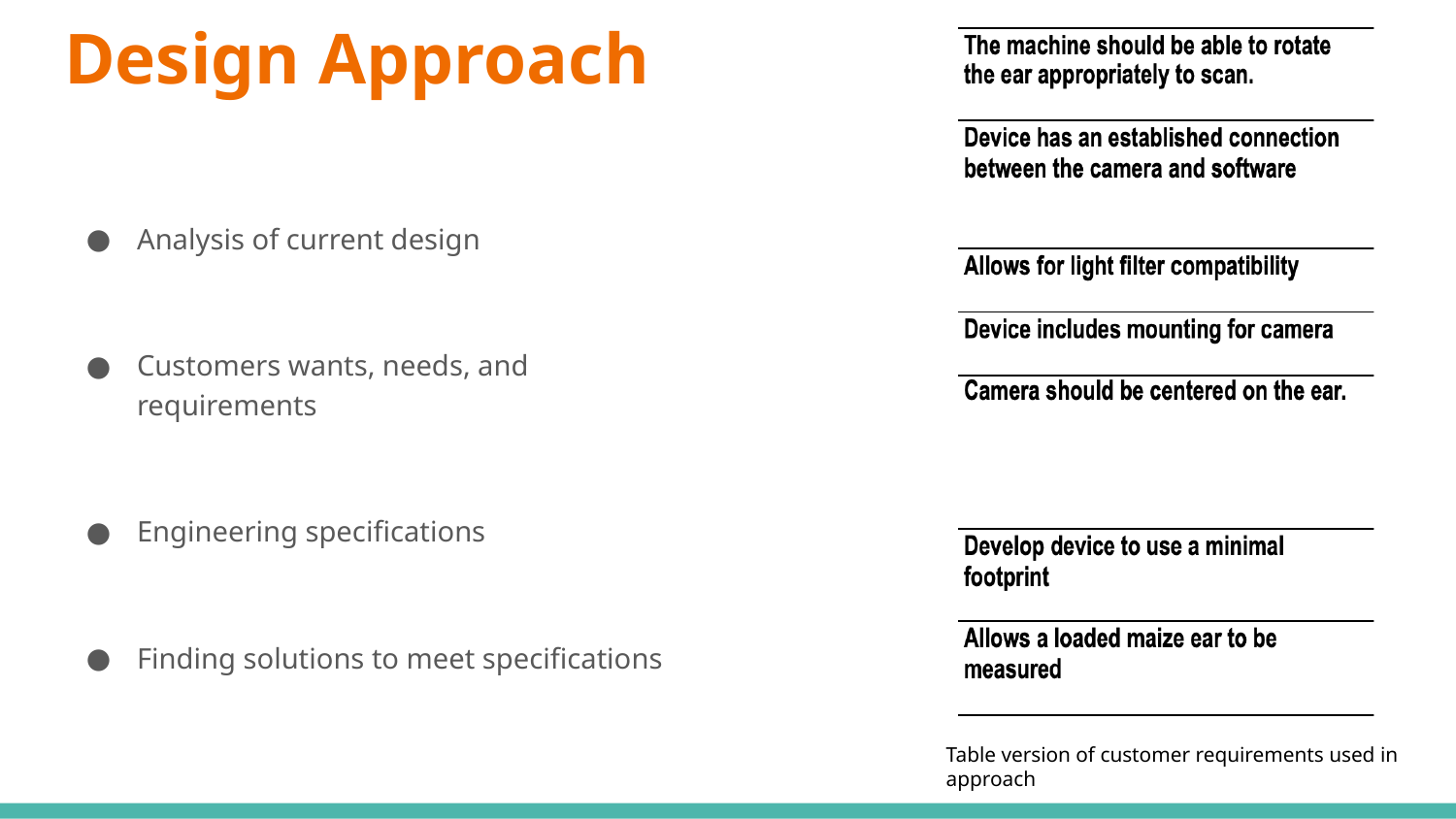

#### Design Approach
Analysis of current design
Customers wants, needs, and requirements
Engineering specifications
Finding solutions to meet specifications
Table version of customer requirements used in approach

### Slide 4
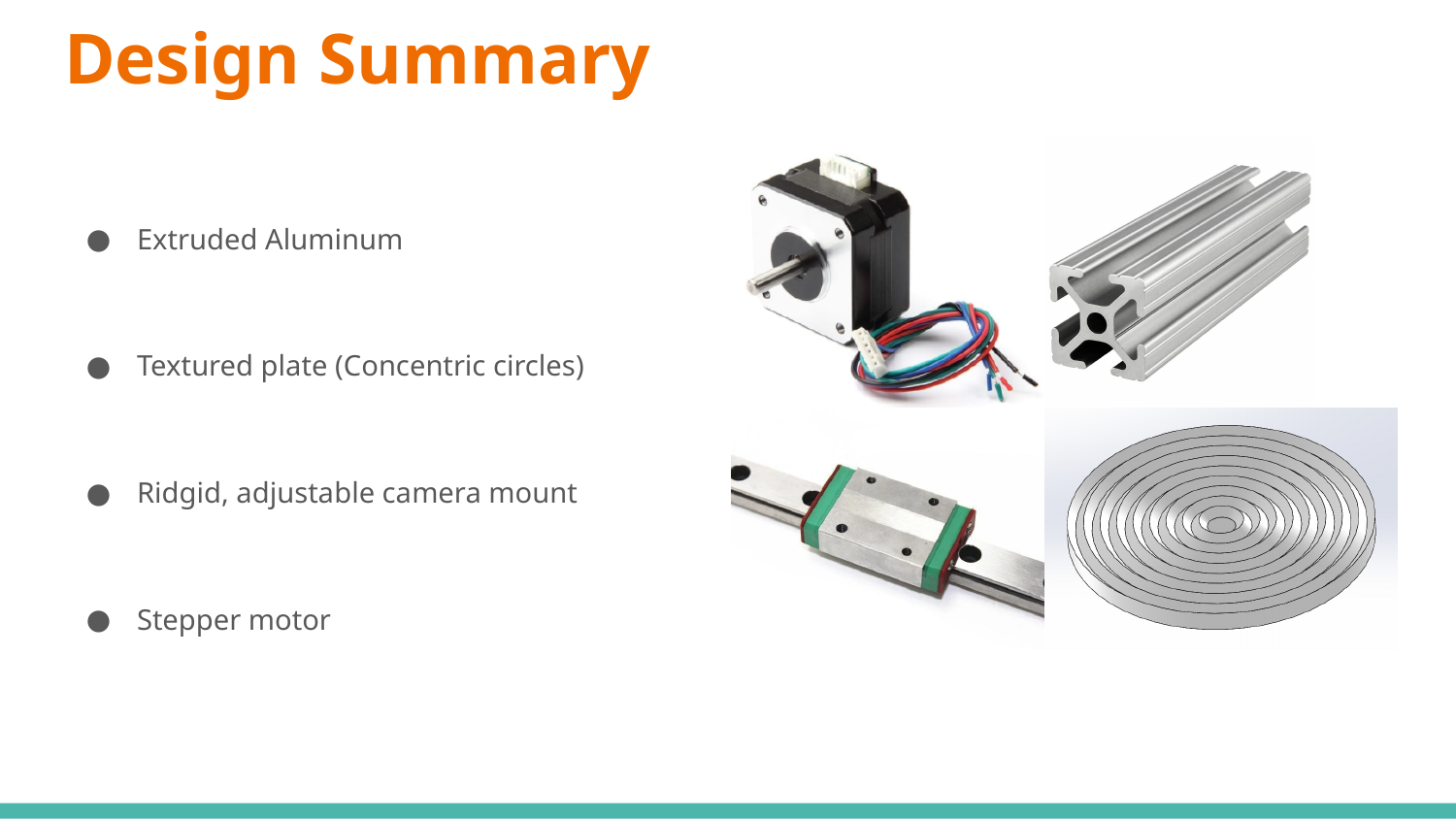

#### Design Summary
Extruded Aluminum
Textured plate (Concentric circles)
Ridgid, adjustable camera mount
Stepper motor

### Slide 5
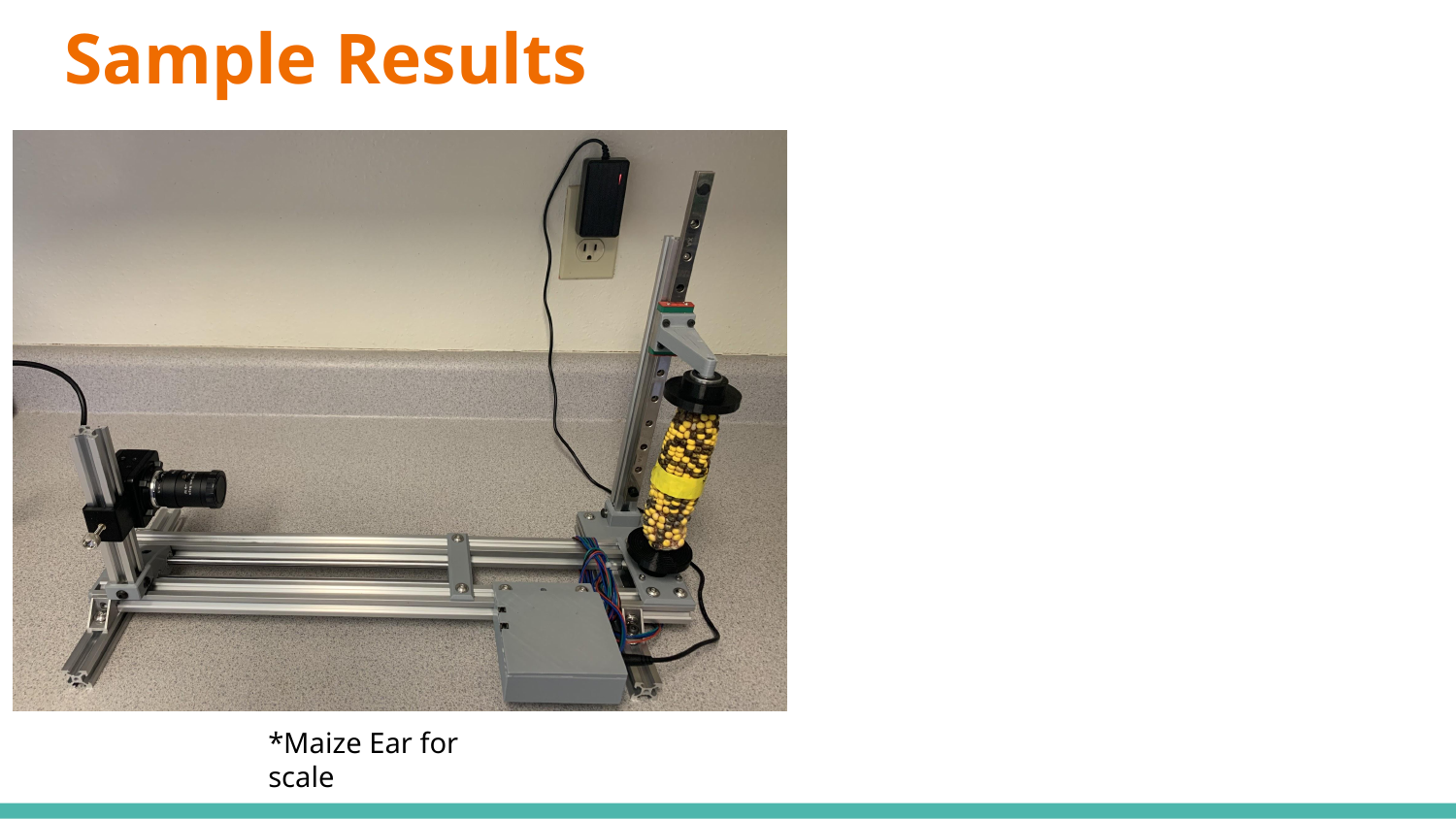

#### Sample Results
*Maize Ear for scale

### Slide 6
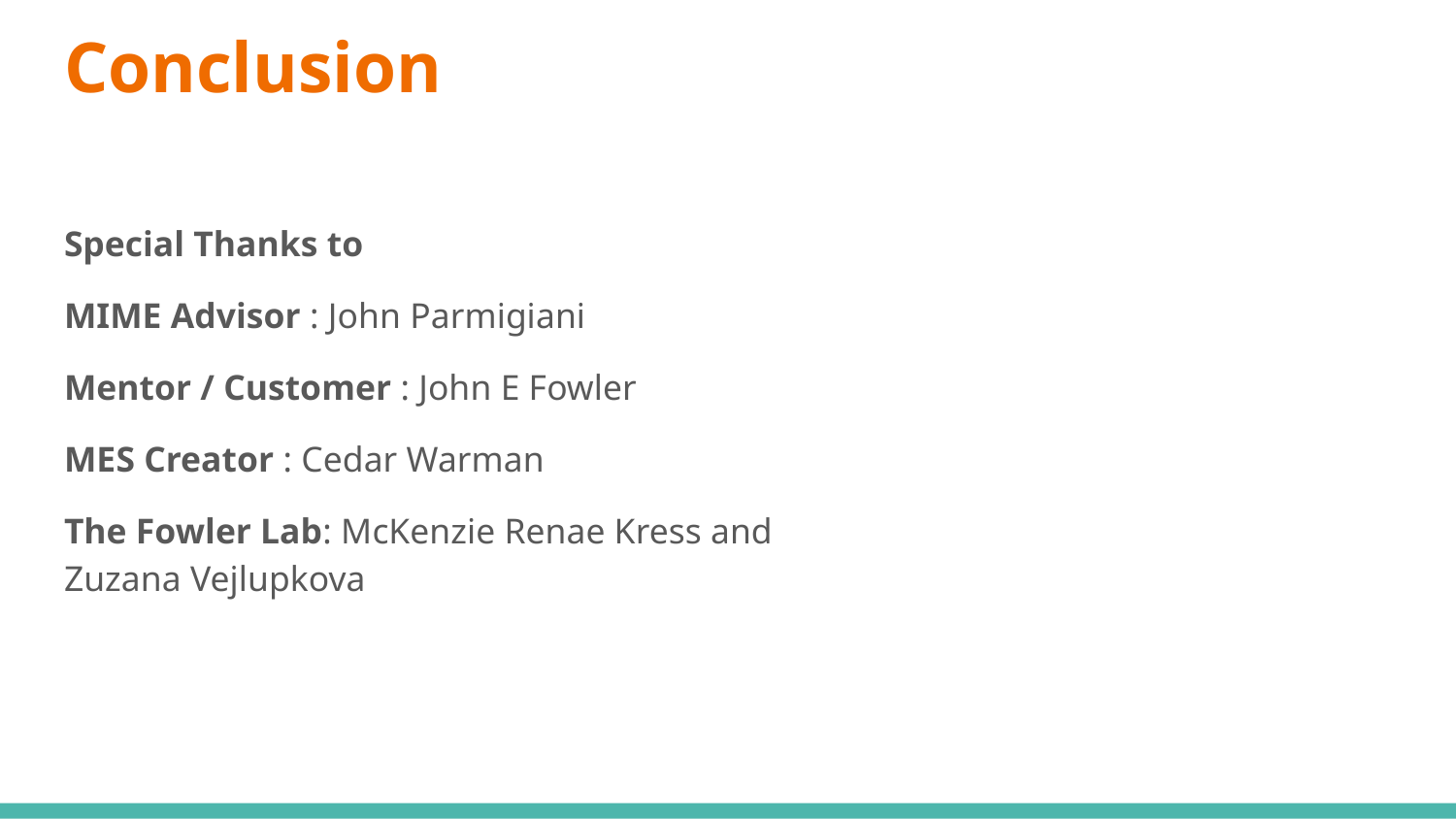

#### Conclusion
Special Thanks to
MIME Advisor : John Parmigiani
Mentor / Customer : John E Fowler
MES Creator : Cedar Warman
The Fowler Lab: McKenzie Renae Kress and Zuzana Vejlupkova
