## Supplementary figures and images for "Spatial inheritance patterns across maize ears are associated with alleles that reduce pollen fitness"

### Schematic_MESv2_2021-07-26.pdf

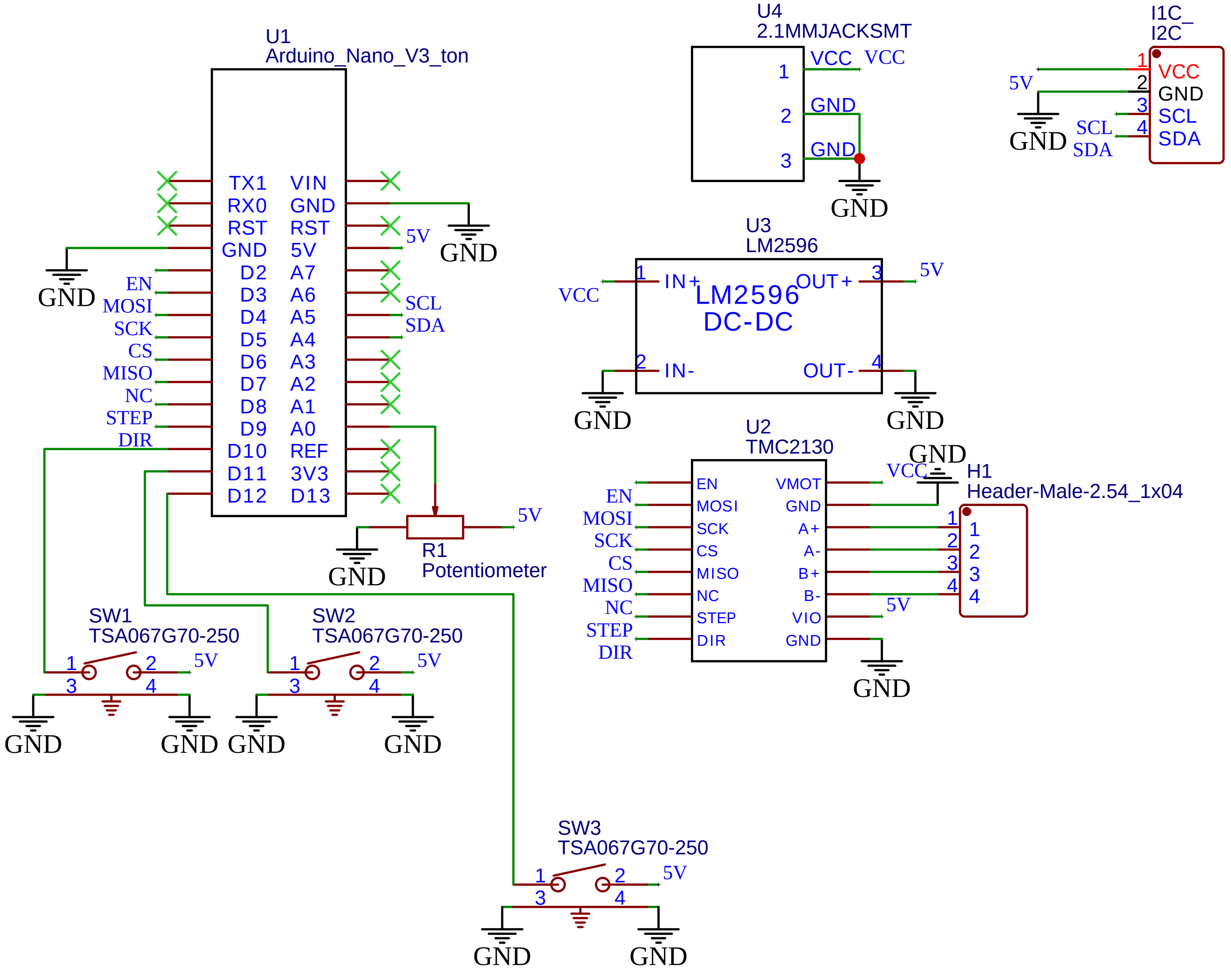


TITLE: Sheet_1

REV: 1.0

Company: Your Company

Sheet: 1/1

Date: 2021-07-26 Drawn By: Taylor Amarotico
