## AppendixS2_Computer_Vision_Development_Supplement for "Spatial inheritance patterns across maize ears are associated with alleles that reduce pollen fitness"

### Overview

The goal of EarVision 2.0 is to develop an accurate but generalized model that could be used to assess images from any field year or dataset. During inference, image assessment includes identification of fluorescent, nonfluorescent, and ambiguous kernels as well as calculation of metrics comparative to ground truth data. These metrics are calculated in order to assess how well the model makes predictions compared to known data.

The first step of this process was to train a new model, adding new images from a recent field season to the training dataset. Initially, EarVision was developed using images from the X and Y datasets (harvested 2018 and 2019, respectively). Since then, the Maize Ear Scanner v2.0 has been developed to image the ears and generate image projections for inference; this includes a new camera and lighting. Images from the B dataset (harvested 2022) thus are associated with differing image parameters from X and Y ear images. Thus, it was important to include some of these in the training dataset in order to retrain the model to be generalizable to a wider range of images.

Next, a series of model retrainings were conducted using this expanded training dataset. New strategies were attempted in order to optimize the results, including a hyperparameter search, image augmentation during training, and identification of potentially problematic images for manual assessment. Most model trainings were performed initially on image projections from B year in order to expedite the process; results that looked promising from that process were used to inform model training using all of the available training data.

Using the best models from the previous step, inferences were run on X, Y, and B datasets separately. The X and Y datasets were the same as those used by Warman et al. 2021, in order to directly compare model results. Finally, inference performance was assessed compared to the ground truth data. The final model applied to EarVision.v2 was one that worked well with all three datasets individually.

### Retraining the Model

#### Model Framework

We utilized Faster R-CNN with a ResNet-50 FPN backbone for training the models, implemented with the Pytorch library. Each model was trained for thirty epochs. Thirty epochs was set as the limit as after that point, there were no longer significant improvements to the models developed.

#### Supporting Libraries

EarVision 1.0 was initially developed using TensorFlow to implement Faster R-CNN. For this version, we switched from using TensorFlow to PyTorch. PyTorch is considered more intuitive and is commonly used in research applications. It abstracts machine learning tasks, reducing the need for low-level coding. In addition, extensive documentation is available specifically for Faster R-CNN with PyTorch, and there is plenty of supporting material available online for reference.

##### How Model Training Works

The model was trained on manually-annotated training data (images and metadata files) generated using Label Studio. The images were randomly split into two sets, the training data and the validation data. Training images were fed into the neural network, with the layers initialized to random weights. The neural network was then applied to the validation set, and compared its results to the manually-annotated metadata to see how well it did. Weights were adjusted accordingly, and the model continues to the next iteration, or epoch. EarVision 2.0 training occurs for 30 epochs. The best epoch for this model is chosen based on the metrics calculated internally, notably the F1 scores and the difference in transmission rate associated each ear, between the values calculated based on the manual validation and model outcomes (see below for details).

##### Parameter Descriptions

Parameters were chosen based off of default values and those we found useful from the hyperparameter search. For default values, 20% of the training set was set aside for validation, the base learning rate was set at 0.005, the number of region proposals to retain for both training and inference was set to 3000 at every stage, with 512 set as the number of region proposals sampled, and IOU threshold was set to 0.7.

Three hyperparameters were tested with the hyperparameter search: number of trainable backbone layers, non-maximum suppression threshold, box score threshold. We found that 4 trainable backbone layers, a non-maximum suppression threshold of 0.5, and a box score threshold of 0.2 worked well for our data and helped to build a reliable and sufficiently accurate model.

#### Training Data

The initial training set contained 300 images from the 2018 (X) and 2019 (Y) field seasons. Manually annotated ears from the 2022 (B) season were added, with a total training dataset size of 409 images. The B images were taken with a new imaging platform (MES.v2), so lighting and image quality are different from X and Y ears. Training images were annotated using Label Studio. In annotation, bounding boxes were drawn around kernels and identified by the person making the annotation as fluorescent, nonfluorescent, or ambiguous. Label Studio saved this annotation data in xml formatted files, with box dimensions given as pixel positions of y_minimum, y_maximum, x_minimum, and x_maximum. Ear images and annotations for the training set is available that the Fowler lab github: https://github.com/fowler-lab-osu

For each training, the data were randomly split into two groups, 20% of total and 80% of total. The 80% was used as training data while the rest was used as validation data. This split would remain consistent throughout the 30 epochs of training for a model, but would be randomly re-split if the model was trained again.

Multiple models were trained on the images with varying strategies, including a hyperparameter search. Images that the model continually had issues with were separated, manually annotated, and introduced to the training set. B-year specific models were also developed and used in inference on the dataset; while these performed well with the B ears, they did not generalize as well as the models developed using the full training dataset (data not shown).

#### Training Set Image Augmentations

As part of the model training process, images were randomly selected for augmentation. Training set image augmentation helps make the model more generalizable, and decreases the likelihood of the model overfitting the data. Augmentations were performed using the Albumentations library. Images had a 50% chance of being flipped horizontally, vertically, or both (Flip); a 20% chance of brightness adjustment by +/- 20% (ColorJitter); a 20% chance of blurring the image (MedianBlur, blur_limit set to 5 to 9); and a 20% chance of a four-point perspective transformation being applied (Perspective, scale set at default values).

#### Average Transmission Difference and Average Absolute Transmission Difference

Metrics were calculated for each training epoch based on the validation (outsample) and training (insample) data separately. The metrics that mostly strongly drove the decision making for the models were the outsample transmission rate difference and the absolute outsample transmission rate difference. Transmission rate is calculated as the percent fluorescent kernels detected out of the total, and is used as a measure of pollen fitness, which is relevant to gametophytic genotype under investigation in each ear. Each ear is a result of two competing pollen genotypes: wild-type (nonfluorescent) vs the mutant allele linked to the fluorescent kernel phenotype, caused by insertion of an engineered *Ds-GFP* transposable element (Warman et al. 2020). The outsample transmission difference (transDiff, Fig. 1) is the difference in the model-predicted value and the known transmission from the validation set, with the positive or negative sign indicating if the value is lower or higher. This is averaged across all images in the validation set for that epoch. Absolute outsample transmission difference (absTransDiff, Fig. 1) is reflective of the magnitude by which this transmission difference occurs. It is worth noting the relationship between these two values: for any model that has a low average absolute outsample transmission difference, a low transmission difference will also occur. However a low average outsample transmission difference does not guarantee a low average absolute outsample transmission difference, as direction of the differences may cancel each other out.

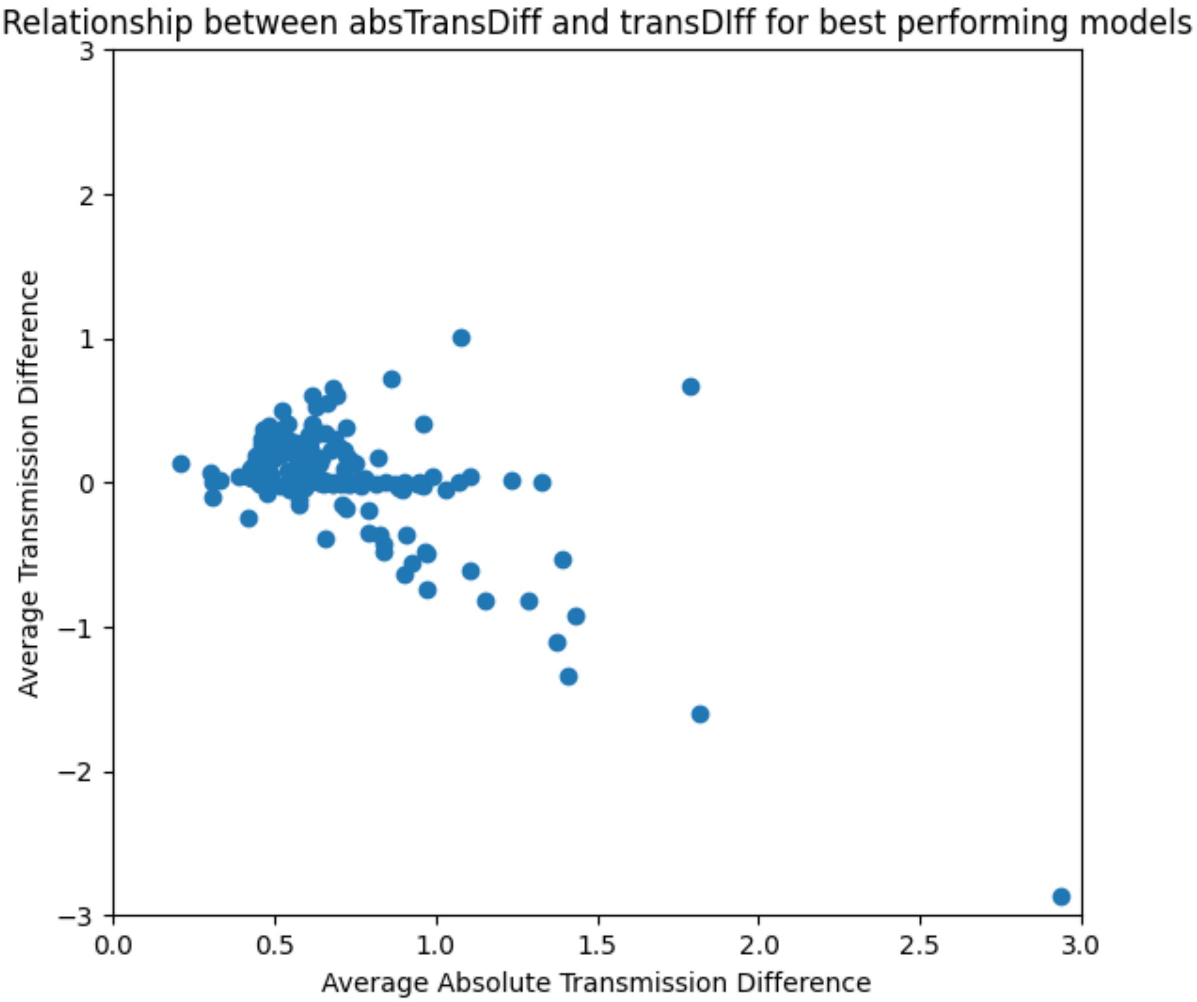


Both values were considered important in selecting the overall best performing model because while the absolute average indicated an accurate model, a certain degree of inaccuracy was considered acceptable if the overall performance of the model was accurate enough.

#### F1 Score Metric

F1 = 2 * TruePositive / (2 * TruePositive + FalsePositive + FalseNegative)

Traditionally, mean average precision is used as the preferred metric for assessing machine learning models. We found, however, that it was extremely time-intensive to calculate mean average precision during training, to the point that other options needed to be considered. We selected the F1 score for assessment to account for both precision and recall. F1 scores for each kernel class (fluorescent and nonfluorescent) were calculated separately, as identification of those classes occurred independently of each other internally.

F1 scores are a measure of the accuracy of the model, accounting for both precision (of all the positive instances of a class, what proportion were true positives) and recall (of all the ground-truth instances of a class, what proportion were accurately identified as true positives). As it is a relationship between two ratios, the ideal value that represents a well-performing model will be close to one.

In order to calculate the F1 score, the model-derived bounding boxes from each class were compared to the ground truth annotations for each image. An IOU threshold of 0.7 and above was considered an acceptable match. The calculation of F1 requires a count of true positive, false positive, and false negative results over the entire image. A true positive is a kernel correctly identified by the model, a false positive was the model finding a kernel where there was none (or identifying a kernel as the wrong class), and a false negative is the model missing a kernel. Each image received its own F1 score for each kernel class. These scores were then averaged across all images but within their own class, yielding two F1 scores for each epoch in the model. In general, we found that F1 scores tended to be higher for fluorescent classification than for non-fluorescent classification, suggesting that EarVision was more successful at positively identifying fluorescent kernels than it was for nonfluorescent. However, the scores for the two different classes tended to correlate(?) to each other, so if a model's F1 for one classification was high, then we could expect its F1 for the other classification to also be high.

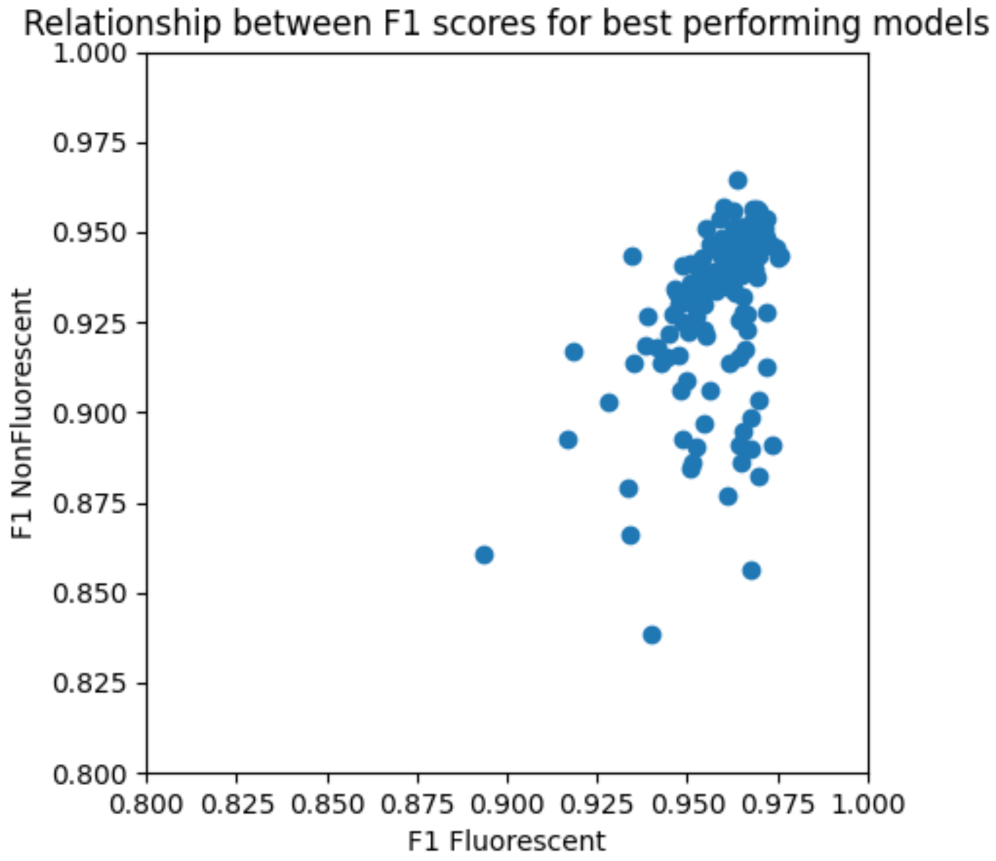


Ultimately we found that higher F1 scores tended to correlate with lower transmission difference and absolute transmission difference metrics. They were thus considered useful for model development, but ultimately less relevant than the metrics based on transmission. F1 served as a useful proxy for comparing the models trained in the earlier iterations of application development as it was faster to calculate; however, mAP was also calculated, for comparison to the initial version of EarVision (Warman et al. 2021).

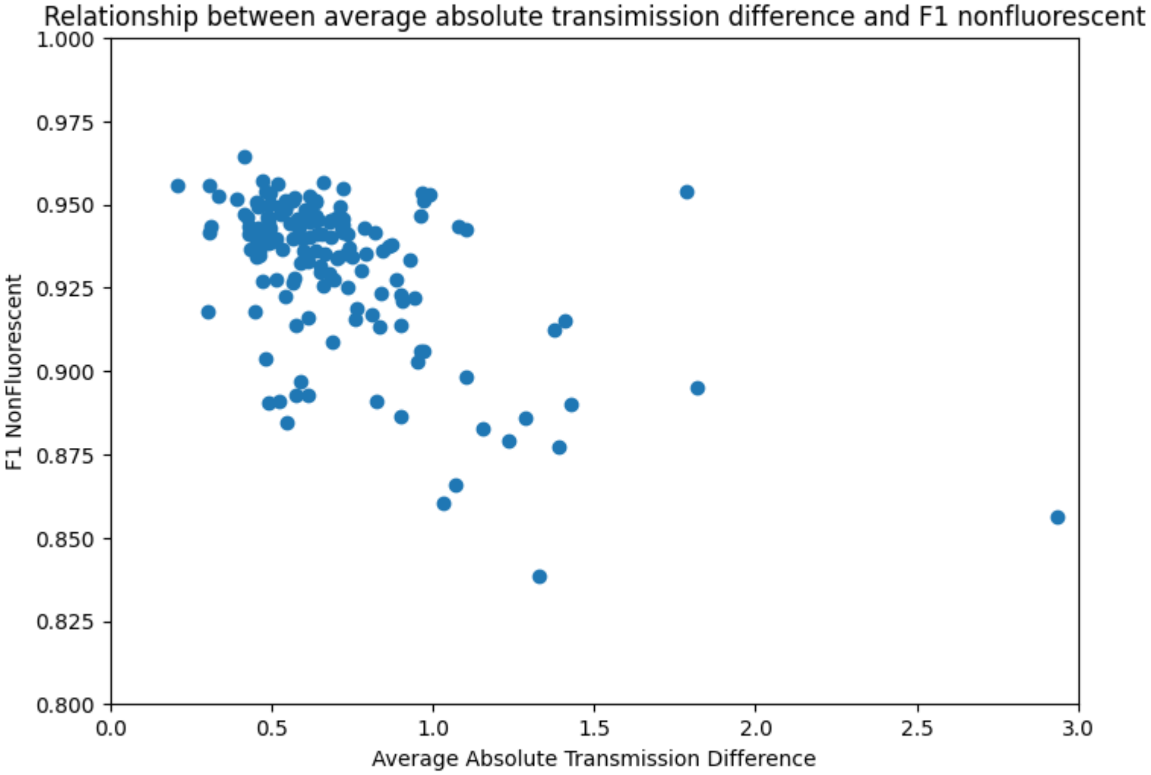


#### Other Metrics

During model development, other metrics were calculated across both insample and outsample results. We chose not to focus on these as outsample transmission metrics and F1 scores represented our preferences for the model performance (overall counts, accuracy of identifying kernels).

#### Hyperparameter Search:

A major strategy in optimizing machine learning models is the hyperparameter search. This is an iterative process that involves testing multiple combinations of hyperparameters and comparing the results. There are a huge number of hyperparameter combinations that can be created for new models, so in the interest of conserving time and effort we chose to focus on three specific hyperparameters that appeared to have the most significant effects on model performance in our early tests: trainable_backbone_layers, box_nms_thresh, and box_score_thresh. trainable_backbone_layers is the number of layers in the neural network that can update their weights during training; box_nms_thresh is the IOU threshold for whether two bounding boxes are identifying the same or different objects; and box_score_thresh is minimum confidence score required for a bounding box to be considered valid.

The search was performed using a grid search strategy. Grid search iterates over all possible combinations of the hyperparameters one would like to investigate; in our search, we completed 150 trainings for 5 possible values of box_nms_thresh and box_score_thresh each, and 6 possible values for trainable_backbone_layers. In the interest of time, mean average precision was not calculated as part of the metrics and only the B 2022 training data was used for training and validation.

The best models that came out of this process were then used to set parameters for the next iteration of test models, these developed using the full training dataset. A smaller, secondary hyperparameter search was completed using the full dataset, varying only trainable_backbone_layers.

Results from the hyperparameter search and subsequent inferences helped us to validate our choice to use the number of ambiguous kernels specifically as a filter for potential hand annotation. A high box_score_thresh value (~0.8) gave models whose inferences had very few to no ambiguous kernels; however, the transmission difference value as well as visual analysis of a few of the images showed that this was due to the model vastly underpredicting kernels of both types. Thus, we chose a lower threshold value, as we saw that overall performance suffered when such underprediction occurred. Rather, we allowed the final chosen model to identify a kernel as ambiguous, and instituted a ‘flag’ once a certain number of those were reached for an image, indicating that those particular images should undergo manual review (see Results in the main text).

Ultimately the hyperparameters chosen as best for overall model performance were: 4 for trainable_backbone_layers, 0.5 for box_nms_thresh, and 0.2 for box_score_thresh. Overall, we found that the hyperparameter search also helped us focus on how to identify problematic ear images, and which metrics were most useful for improving the inference process.

#### Comparing Inferences

##### General Outcome

In general, different models performed well with different datasets, even when the models were trained on the full training dataset (409 images containing X, Y, and B ears). Of four final, well-performing models assessed, the model dubbed “José” was the most generalizable, performing well with all three datasets.

Given that two of these datasets were the same used in Warman et al 2021, with their performance assessed with the initial, year-specific EarVision models in Figure 5, we calculated similar values to compare the new José model with the year-specific models described in that paper. We found that José performed similarly to the single year models, but could generalize across images from all three field years (and three different digital cameras/versions of the Ear Scanner). Note that these performance metrics do not include the implementation of the ambiguous kernel ‘flag’ function discussed in the main text (Figure 3, Supplemental Figures 2 and 3).

| Dataset | Warman Fluorescent Rsq | José Fluorescent Rsq | Warman Non-Fluorescent Rsq | José Non-Fluorescent Rsq | Warman Transmission Rsq | José Transmission Rsq |
|---|---|---|---|---|---|---|
| X 2018 | 0.986 | 0.989 | 0.977 | 0.990 | 0.984 | 0.955 |
| Y 2019 | 0.984 | 0.992 | 0.969 | 0.911 | 0.945 | 0.935 |
| B 2022 | n/a | 0.986 | n/a | 0.983 | n/a | 0.978 |
